# Optimizing short-channel regression in fNIRS: an empirical evaluation with ecological audiovisual stimuli

**DOI:** 10.1101/2025.05.23.655796

**Authors:** Y. Lemaire, P. Barone, K. Strelnikov, O. Deguine, A. Caclin, J. Ginzburg

## Abstract

**Significance:** Functional Near-Infrared Spectroscopy (fNIRS) is increasingly favored for its portability and suitability for ecological paradigms, yet methodological standardization remains a challenge regarding the optimal use of short-separation channels (SC) to remove systemic physiological noise.

**Aim:** We aim to evaluate and compare methods of SC regression as implemented in the most widely used fNIRS analysis toolboxes, with the goal of reaching a consensus on best practices for incorporating SC into generalized linear model (GLM)-based analyses. Specifically, we compared ten SC regression strategies addressing SC selectivity, dimensionality reduction strategies, and SC availability.

**Approach:** 16 healthy adults passively listened and watched ecological auditory, visual, and audiovisual stimuli while occipital and bilateral auditory cortices were recorded.

**Results:** Oxygenated hemoglobin signals (HbO) processed without SC regression produced uninterpretable results in the context of the present study. Non-selective SC regression methods that pooled all available SC signals consistently outperformed anatomically or functionally restricted approaches. Orthogonalization further enhanced performance by reducing redundancy and capturing shared systemic variance, improving detection of stimulus-specific cortical responses and contrast sensitivity. Deoxygenated hemoglobin signals (HbR), while less sensitive to systemic artifacts than HbO, benefited most, similarly to HbO, from the pooled, orthogonalized SC signal approach.

**Conclusion:** Overall, our findings highlight the essential role of SC regression in recovering physiologically meaningful signals in fNIRS and recommend including all available SC channels within the GLM, coupled with orthogonalization techniques, as a generalizable best practice for denoising across hardware configurations.

## 1. <u>Introduction</u>

Functional Near-Infrared Spectroscopy (fNIRS) is an increasingly prominent neuroimaging modality, valued for its portability, affordability, and non-invasiveness^1^. By measuring relative changes in oxy- (HbO) and deoxy- (HbR) hemoglobin concentrations in cortical tissue, fNIRS enables the investigation of a wide range of cognitive functions, including perception, attention, memory, language, and motor processing^2^. Its ease of use and tolerance for motion makes it particularly well suited for studying populations traditionally underserved by neuroimaging research, such as infants, children, older adults, and clinical groups. Importantly, fNIRS is especially promising in the field of auditory neuroscience. Unlike functional Magnetic Resonance Imaging (fMRI), it is silent—an essential feature when studying auditory processing. Compared to electroencephalography (EEG) and magnetoencephalography (MEG), fNIRS is unaffected by hearing aids or cochlear implants, offering a unique advantage for studying individuals with hearing impairments. As such, fNIRS has been used in a growing body of research on auditory perception, including investigations into the processing of physical features of sound like intensity levels or auditory-spatial cues^3–6^, as well as research on auditory cognitive processes like speech processing, listening effort, language development, development of auditory cortical function, and auditory short-term/working memory^3,7–9^. It has also shown promise for clinical applications such as in the study of cortical reorganisation after impaired sensory input or in tinnitus and during subsequent rehabilitation with cochlear implants^8,10^.

Yet despite these many advantages, several key methodological challenges still limit the full potential of fNIRS. One major challenge lies in the diversity of preprocessing practices across research groups and toolboxes, which makes it difficult to compare findings and reach generalizable conclusions^1^. A central issue in fNIRS data analysis concerns the optimal use of short separation channels (SCs) for removing systemic physiological noise; while it is well established that incorporating SCs improves signal quality compared to not using them, there remains no consensus on which SC-based approach is most effective. The present study aims to address this issue by systematically evaluating different strategies to use SC information for systemic signal removal, in the context of passive, ecologically valid auditory, visual, and audiovisual stimulation, and across cortical regions.

### 1.1. Ecological stimulation: an underused opportunity

A growing body of neuroscience research calls for using ecological stimuli to better approximate real-world perception^11^. In this regard, fNIRS is uniquely positioned to bridge the gap between laboratory experiments and everyday cognition, given its robustness to motion and its flexibility across settings^12^. However, few fNIRS studies have taken full advantage of this by incorporating complex, real-life stimuli such as movies or audio stories. Despite this potential, to our knowledge, no study has yet systematically examined the effectiveness of different analysis strategies when applied to fNIRS data acquired during passive, ecological stimulation involving auditory, visual, and audiovisual content. In the current study, while we employed a structured block design, we exclusively used video clips and sounds of real-life objects and events, thus defining them as ecological, in contrast to paradigms that rely on more artificial stimuli (e.g., pure tones, geometric shapes), as an effort to approximate more realistic stimulation conditions.

### 1.2. Challenges in signal quality and systemic noise

While fNIRS is praised for its practical advantages, it is not without limitations. One of the main challenges lies in the contamination of neural signals by systemic physiological noise including heartbeat, respiration, low-frequency blood pressure oscillations, Mayer waves, and spontaneous neural activity^13–16^. These confounds can originate from both superficial extracerebral layers and deeper cortical tissues^17^, and can significantly obscure the measurement of true cerebral responses. While certain components of systemic physiological noise (i,e., low-frequency blood pressure oscillations, heart rate, and respiratory signals) fall outside the frequency range of the hemodynamic response and can be effectively attenuated using temporal filtering (e.g., band-pass filtering^18^), this approach has clear limitations. In particular, it cannot eliminate low-frequency oscillations in blood flow, such as Mayer waves (∼0.05 to 0.15 Hz), which overlap spectrally with the expected signal in block design paradigms^14,16,19^. These slow vascular fluctuations, driven by systemic physiology rather than localized neural activity, remain a major confound for interpreting fNIRS data and need targeted methods for removal.

### 1.3. The unresolved issue of short-channels in fNIRS

SCs are a powerful tool for isolating superficial hemodynamic signals, particularly those arising from extracerebral systemic activity such as cardiac, respiratory, or Mayer waves. By positioning detectors at short distances from the source (typically ∼8 mm), SCs primarily capture these superficial signals, which can then be regressed from the long-distance channel signals (that measure both the cerebral and superficial signals) to improve the specificity of the cortical response. Before SCs became widely available from most fNIRS manufacturers, researchers relied on alternative methods to mitigate systemic contamination (e.g., independent component analysis^20^; HbO-HbR-correlation based correction^21,22^). While these methods provided partial solutions, the introduction of SCs marked a significant advance by offering a direct, empirical estimate of extracerebral physiology, an approach that has been shown to outperform other correction methods in terms of removing systemic artifacts and improving signal specificity^23–28^.

Despite growing recognition of their importance, the use and implementation of SCs in fNIRS studies remain inconsistent. In a large multi-lab collaborative study, Yücel et al.^1^ brought together 102 researchers from 38 international teams to analyze two fNIRS datasets. Their findings revealed a striking disparity in analysis pipelines and outcomes, underscoring the lack of consensus within the field and the urgent need for standardized preprocessing approaches. Specifically, 79% of the teams adopted a generalized linear model (GLM) framework, which enables the inclusion of SC signals as regressors. Among these, 46% used individual SC signals as single regressors, 6% used the first principal component of all SC signals, 39% applied a multiple-regressor approach using either raw SC signals or their principal components. For the remaining 9%, the methodology could not be determined from the available information. However, none of the teams reported the number or placement of SCs used, nor did they justify the rationale behind their chosen implementation.

This lack of consensus is also reflected in the divergent implementations offered by the main fNIRS signal processing toolboxes. Although including SC regressors in a GLM framework is now considered standard practice, the specific method for doing so varies substantially^29^. For instance, Homer3 primarily implements a spatially-specific correction, where the signal from the nearest SC is regressed from each long channel on a one-to-one basis^30^. Homer3 also implements a correction where the signal from the most correlated SC is regressed from each long channel, also on a one-to-one basis. In contrast, MNE-NIRS primarily recommends a global correction approach, such as using the average of all SC signals as a single systemic regressor^23^. The NIRS Brain AnalyzIR Toolbox, by default, performs a global correction approach, implementing regression using all components derived from a principal component analysis (PCA) of the signals from all SCs for both HbO and HbR^31^.The co-existence of these fundamentally different strategies (local vs global, PCA-based or not) mirrors the findings of Yücel et al.^1^ and highlights the methodological variability that motivates the present study, which directly compares these approaches.

When using SCs, two main scenarios typically arise: (1) each source optode is paired with a nearby detector to form a short-separation channel, or (2) only a subset of source optodes are used to create short-separation channels, due to hardware constraints. Within each of these scenarios, multiple strategies exist for regressing SCs from the long-channel signals. One approach favors spatial specificity, regressing each long channel with the SC signal nearest to its source-detector pair^30^. A related signal-specific approach regresses each long channel with the SC signal to which it is most strongly correlated^30^. In contrast, non-selective approaches either regress the average of all raw SC signals (computed separately for HbO and HbR) from all long channels^23^, or use all available SCs to extract global components (e.g., via principal component analysis, PCA) and regress these from the long channels^31^, thus targeting systemic noise that is global in nature but less spatially precise. This question becomes especially important in multi-lobar studies, where systemic interference might differ across anatomical regions, such as between temporal and occipital lobes, and decisions about SC selectivity versus generality in SC regression could critically impact the interpretation of results.

The co-existence of these fundamentally different strategies highlights a methodological ambiguity. Given that decisions about SC selectivity versus generality in SC regression could impact the interpretation of results, this lack of consensus underscores the need for standardized preprocessing approaches. Establishing a generalizable best practice for SC regression is therefore essential for recovering physiologically meaningful signals and improving the comparability of fNIRS findings across studies, and directly motivates the present work.

### 1.4. Objectives of the current study

The current study aims to systematically address the unresolved questions surrounding the use of SCs in fNIRS preprocessing, with a particular focus on their role in block-design GLM analyses. We recorded fNIRS signals over the temporal and occipital lobes while administering passive, ecological auditory, visual, and audiovisual stimuli to healthy participants. We formulated clear a priori hypotheses regarding expected brain responses (i.e., occipital channels for visual conditions and temporal channels for auditory conditions). This approach is grounded both in decades of fMRI research and in recent fNIRS studies showing that dynamic visual and auditory stimuli reliably elicit activation in these regions^5,32–38^. Optode placement followed a well-established, atlas-based selection procedure commonly used in fNIRS research^24,39,40^. To evaluate pipeline performance, we implemented two complementary metrics. The first quantified the emergence of activity, compared to a no-stimulation baseline, in each channel that was specific to the task and sensory modality, indicating whether expected activation could be detected at the individual-channel level. The second metric assessed differences between conditions within these channels to ensure that the observed effects followed the predicted pattern of responses, providing an additional check against spurious activations. This approach, consistent with previous studies on SC regression in fNIRS data analysis^23,24,40^, assumes that effective preprocessing should reveal activation in channels located over the expected functional regions. Counting such emergent channels thus provides an empirically grounded approximation of pipeline performance, while the second metric serves as a complementary, functionally constrained validation of these effects. Importantly, our montage was designed to include one SC per source, allowing us to explore a variety of SC correction strategies within the same dataset. We tested ten distinct preprocessing pipelines:

1. No SCs (baseline condition)
2. Limited SC availability (n = 3):

a. spatial-specific approach (nearest)
b. signal-specific approach (most correlated)
c. non-selective mean-based approach
d. non-selective approach with PCA
3. Full SC availability (n = 8):

a. spatial-specific approach (nearest)
b. signal-specific approach (most correlated)
c. non-selective mean-based approach
d. non-selective approach without PCA
e. non-selective approach with PCA

## 2. METHODS

### 2.1. Participants

Sixteen participants, aged 24 to 34 years (M = 27.1, SD = 2.7; 8 females), were recruited for this study. All participants self-reported having no auditory or uncorrected visual impairments, and none of the participants had any history of neurological or psychiatric conditions, nor had they been diagnosed with any neurodevelopmental disorders. The study was approved by the OUEST IV ethics committee (approval number: 2019-A00295-52) and was conducted in compliance with the Declaration of Helsinki. Informed consent was obtained from all participants before their involvement in the study.

### 2.2. Stimuli

Visual stimuli were presented on a 27” screen (DELL, P2719H) with a 60 Hz refresh rate. Auditory stimuli were delivered via speakers (KAIWIN SP 690N), calibrated to produce a sound level of approximately 60 dB SPL (A weighted). We exclusively used ecologically valid stimuli, comprising video clips or sounds of real-life objects and events (e.g., moving cars, a ringing phone). Fifteen distinct concepts (e.g., ’bird’, ’sheep’, ’boat’) served as the basis for stimulus generation. For each concept, three corresponding stimuli were created: an auditory-only version (e.g., the sound of birds singing presented with a black screen), a visual-only version (e.g., a peaceful meadow), and an audiovisual version (e.g., a video of birds singing in a meadow with the corresponding sound). All stimuli were unique and sourced from royalty-free content websites. Each individual stimulus presentation lasted for 10 seconds.

### 2.3. fNIRS Montage and Data Acquisition

Near-infrared light absorption was measured at 760 nm and 850 nm wavelengths at a sampling frequency of 7.81 Hz using a continuous wave NIRScout instrument (NIRx Medical Technologies, LLC). Data were collected using the NIRStar 15.2. acquisition software. Eight light sources and eight light detectors were attached to a cap with a 10-20 system to the probe location (see Figure 1). In addition, eight short 8-mm channels (one for each source) recorded systemic signals. The montage was created using fOLD (fNIRS Optodes Location Resolution)^41^, which allows placing the optodes in the 10-20 international system to cover selected anatomical areas. We used the Brodmann atlas^42^ to generate a montage covering the auditory and visual cortex. For the auditory cortex, we targeted the left and right Superior Temporal Gyrus (STG) and the Middle Temporal Gyrus (MTG). For the visual cortex, we targeted the Primary Visual Cortex (V1) and the Visual Association Cortex (V2). Occipital channels were treated as a single region of interest (ROI) because they are positioned primarily along the midline or in close symmetry, providing limited lateral separation. In addition, since our main goal was to assess pipeline performance rather than hemispheric differences, dividing the occipital cortex into left and right ROIs would have added complexity without methodological benefit. One channel located over the right posterior temporal area (channel 16) was excluded from the analysis because it did not have a left-hemisphere counterpart. The final montage thus included fifteen measurement channels with source-detector distances ranging from 2.5 cm to 4 cm. In total, 7 channels were covering the occipital area and 4 channels per hemisphere were covering bilateral temporal areas. The detail of the location of each channel and its two highest specificities to Brodmann areas are presented in Table 1.

**Figure 1:**
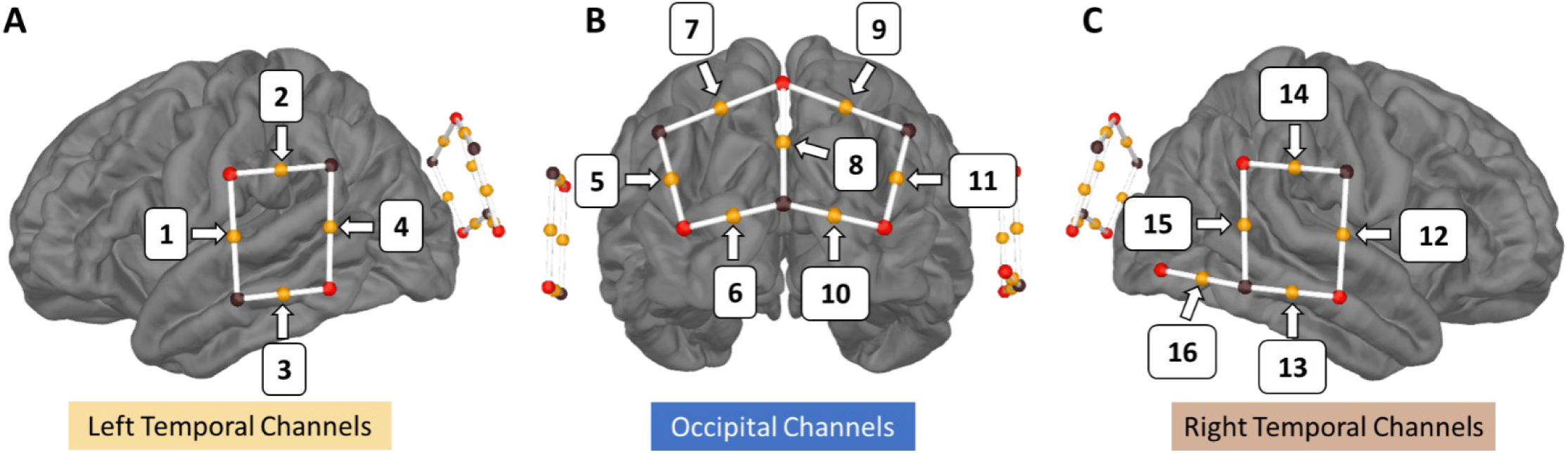
Location of sources (black spheres) and detectors (red spheres) and their corresponding midpoint channels (orange spheres with white lines connecting the source/detector pair for each channel) on (A) left; (B) back; and (C) right views of the brain. Channel 16, located in the left-most part of the right temporal montage, is the only channel without a corresponding counterpart on the left side (see also Table 1). It was excluded from further analysis due to this asymmetry. The visualization was obtained using the mne-nirs library23.

**Table 1:**
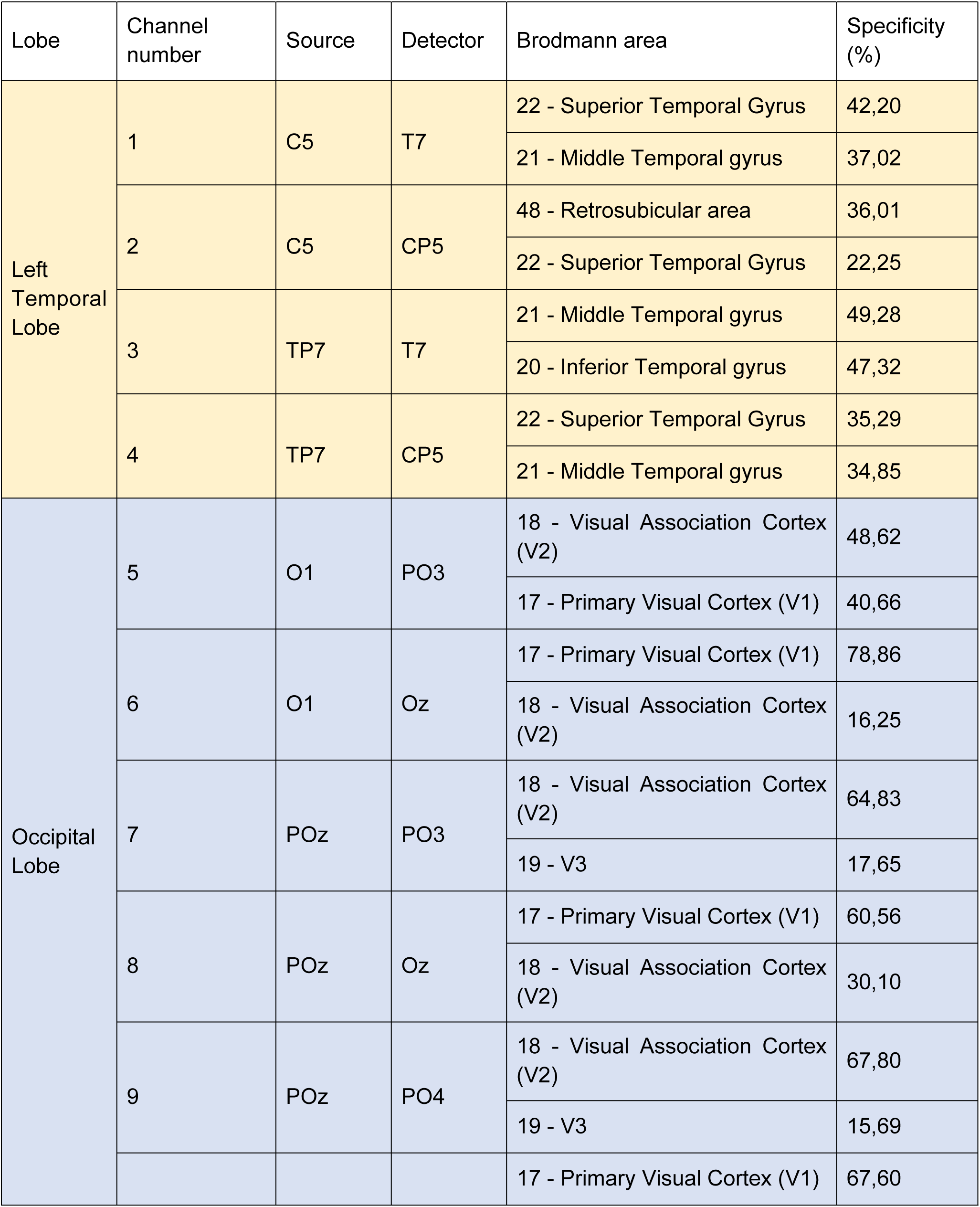
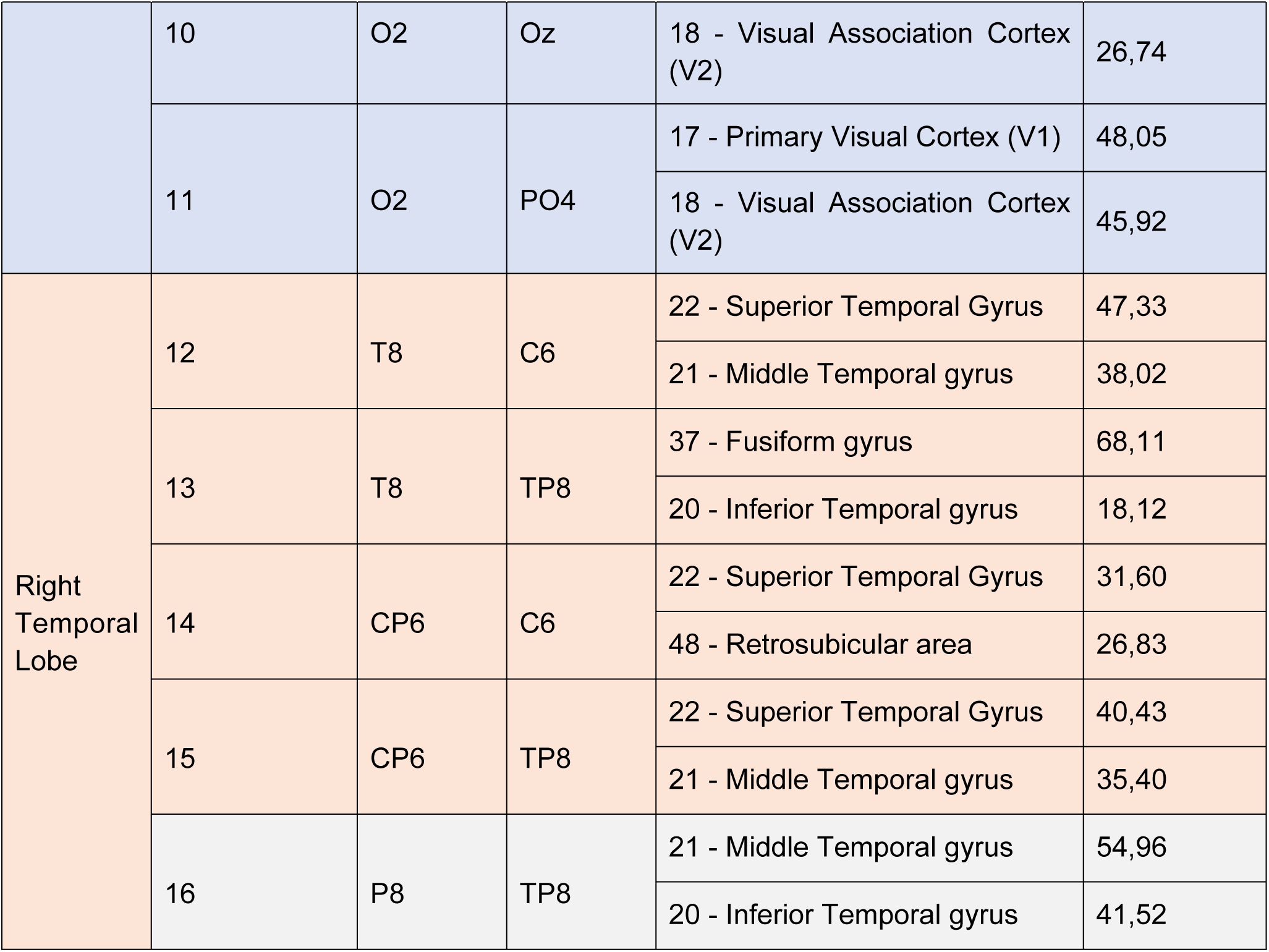
Channel specificity: each of the 16 recording channels is composed of a source and a detector (located in standard 10-20 positions). We reported in the table the Brodmann areas for each channel with the highest specificity (in percent) according to the fOLD software^41^. Channels localized over the occipital lobe are highlighted in light blue. Channels over the left temporal area are highlighted in light yellow, and channels over the right temporal area are highlighted in light red. Channel 16, highlighted in light grey, was excluded from statistical analysis because it did not have a left-hemispheric counterpart.

### 2.4. Procedure

The experimenter measured the participant’s head circumference before the session to establish the proper cap size. The cap alignment was meticulously checked and adjusted to place the Cz optode halfway between the two tragi and between the nasion and inion. The participant was then brought into a dimly lit, sound-attenuated booth. If necessary, their hair was gently moved using a thin wooden stick to allow unobstructed access to the optode locations. Optodes were positioned according to the pre-defined montage (see Figure 1 and Table 1). The fNIRS signal was calibrated and monitored for quality, and optode placement was adjusted as needed until satisfactory signal quality was achieved before continuing. Participants were instructed to limit their movements, avoid talking, listen carefully, and keep their eyes on the screen during the experiment while they were seated comfortably. Participants were given a 60-second baseline period to focus on a central crosshair before the experiment started. Passive listening and viewing only were necessary for the task itself.

Participants completed a continuous sequence of 45 trials, consisting of 15 trials for each condition: auditory-only (A), visual-only (V), and audiovisual (AV). The presentation order was pseudo-randomized, adhering to the constraint that each consecutive group of three trials contained one trial from each sensory modality (A, V, AV), with the specific order within each triplet being random. This pseudo-randomized mini-block design was chosen to minimize habituation effects that can occur in long, single-condition blocks, to maintain participant engagement across modalities, and to preserve trial unpredictability. Following each 10-second stimulus presentation, there was a silent inter-stimulus interval (ISI) with a duration jittered between 8 and 12 seconds. During this ISI, participants were instructed to fixate their gaze on a static black-and-white image of a cartoon planet Earth displayed in the center of the screen. The total duration of the experimental task, including the initial 60-second baseline, ranged from 14.5 to 17.5 minutes.

### 2.5. fNIRS data preprocessing

fNIRS data were preprocessed using the NIRS Brain-AnalyzIR toolbox^31^ along with custom-written scripts on Matlab R2021a (available at https://github.com/yanndavlem/nirs_article). We first converted raw intensity signals to changes in optical density. To mitigate the impact of motion artifacts, which can arise even from small, involuntary movements, we applied Temporal Derivative Distribution Repair (TDDR)^43^. This method was chosen for its selective nature: it identifies artifacts by detecting data points with a statistically improbable rate of change (i.e., sharp spikes) and corrects only these points, leaving the slower, underlying hemodynamic signal of interest unaffected when no artifacts are present. The corrected optical density was then band-pass filtered between 0.01 and 0.12 Hz to remove cardiac activity (∼1.2 Hz) and respiratory activity (∼0.25 Hz). Subsequently, the corrected and filtered optical densities were transformed into (de)oxygenated hemoglobin concentration changes using the modified Beer-Lambert Law, incorporating a fixed pathlength factor of 0.1, which corresponds to the default setting in the NIRS Brain-AnalyzIR toolbox^31^. After visually inspecting the preprocessed data, no data were excluded from subsequent analyses.

### 2.6. Regression of systemic signal with short-channels

We conducted Generalized Linear Model (GLM) analyses to evaluate the impact of SC regression on fNIRS signals (Figure 2). All analyses were performed using Ordinary Least Squares (OLS), a widely used method for estimating the relationship between a dependent variable and one or more independent variables by minimizing the residual sum of squares^44^. This approach allowed us to obtain beta coefficients that quantify the relationship between the experimental conditions and the hemodynamic responses.

**Figure 2:**
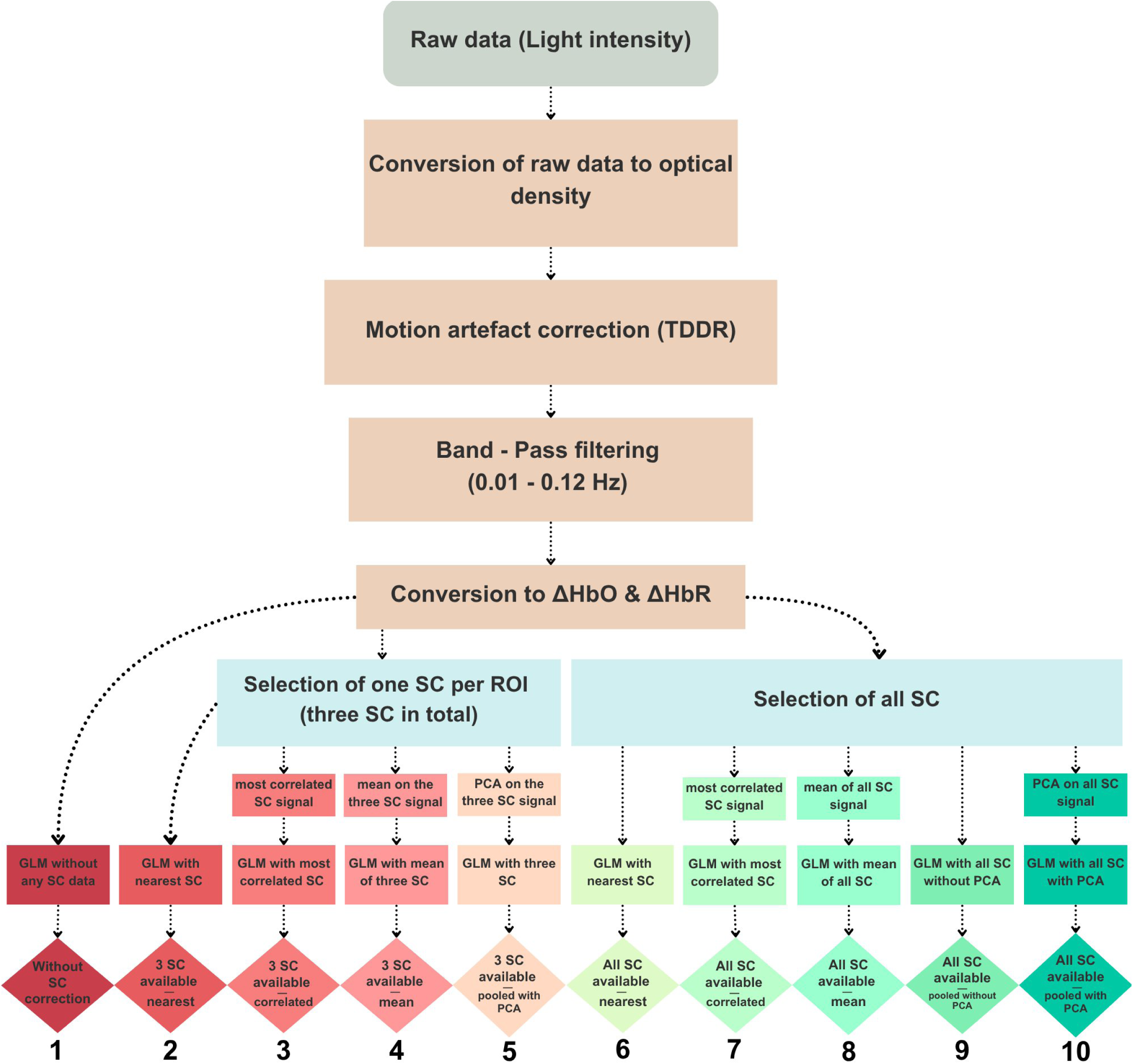
Preprocessing and analysis pipelines of the fNIRS data. A standard pipeline was used to convert raw signals to HbO and HbR concentration changes (ΔHbO, ΔHbR). The main statistical analysis (GLM) was performed using either no SC correction, three SCs (to simulate potential hardware limitations), or all eight available SCs. Four correction approaches were applied: a spatial-specific method, in which each LC was regressed using the signal from the nearest SC, a signal-specific approach in which each LC was regressed using the signal from the most correlated SC, and two non-selective methods, in which SC signals were either averaged or pooled together to regress each LC. In the latter case, we additionally assessed the impact of applying a PCA to the SC signal. This resulted in ten distinct pipelines: (1) No SC correction, (2) 3 SCs – nearest (spatial-specific), (3) 3 SCs – correlated (signal-specific), (4) 3 SCs – mean (non-selective), (5) 3 SCs – pooled with PCA (non-selective), (6) All SCs – nearest (spatial-specific), (7) All SCs – correlated (signal-specific) (8) All SCs – mean (non-selective), (9) All SCs – pooled without PCA (non-selective), and (10) All SCs – pooled with PCA (non-selective). SC: short-channels; LC: long-channels; PCA: principal component analysis; GLM: generalized linear model. TDDR: temporal derivative distribution repair.

The predictor variables were defined using a boxcar function convolved with the canonical Hemodynamic Response Function (HRF), allowing us to generate a Statistical Parametric Mapping of the HRF (SPM-HRF)^31^ for each of the three conditions (A, V, and AV). In all analyses, a GLM was applied to each long-channel and each chromophore (HbO, HbR) with one HRF-convolved regressor for each condition. The same canonical HRF, which models a positive signal deflection, was used for both chromophores; consequently, the expected physiological decrease in HbR is represented by a negative beta value.

In addition to the three regressors for each condition, the number and nature of the regressors varied according to the SC strategy used, ranging from 0 (no SC regression) to up to 16 regressors (when including all available SCs of both chromophores, with or without PCA). To evaluate different strategies for SC correction, we compared two main scenarios: one in which only three SCs were available (to simulate hardware limitations) and another using all eight available SCs (one per source). For the limited SC scenario (n=3), we selected one representative SC per lobe to reflect a pragmatic placement strategy based on montage geometry. For the occipital region (channels 5-11), we selected the SC at source POz because its location on the sagittal midline provides a natural point of symmetry for the bilaterally placed channels within that lobe. For the temporal lobes, no central optode was available in our montage, so we chose one of the two SC sources arbitrarily. The SC at source CP6 was chosen for the right temporal area (channels 12-15), and the SC at source C5 was chosen for the left temporal area (channels 1-4). Within each scenario, we used one of three strategies: a spatial-specific approach, where each long channel was corrected using its nearest SC; a signal-specific approach, where each long channel was corrected using the SC with which it was most correlated; or a non-selective approach. For the non-selective approach, we either used the MNE-NIRS implementation^23^, where each long-channel was corrected using the average of all SC signals, or the NIRS Brain AnalyzIR implementation^31^ where all SC signals were pooled to model widespread physiological interference across the montage. In the latter case, when all SC signals were available, the GLM was run with and without applying PCA to evaluate its impact on capturing global physiological components shared across sources.

For the non-selective approaches employing PCA, the following procedure was applied. The time series from all available SCs for both chromophores were first concatenated into a single matrix (i.e., [time points × 6 signals] for the limited scenario, and [time points × 16 signals] for the full scenario). A PCA was then applied to this combined matrix. All resulting principal components were retained and subsequently included as regressors of no interest in the GLM for each long channel and chromophore, following the NIRS Brain AnalyzIR^31^ toolbox implementation.

Six SC correction strategies were thus tested:

1. No SCs (baseline condition) This reference analysis involved no SC regression. Only standard pre-processing steps and a GLM with the three condition regressors were applied.
2. Limited SC availability (n = 3) For the next three analyses, we simulated scenarios in which only three SCs were available, reflecting potential hardware limitations or experimental constraints. In this configuration, one SC was selected per lobe based on its central location relative to surrounding long channels (LCs): the SC near source C5 was used for the left temporal region (channels 1–4), the SC at POz for the occipital region (channels 5–11), and the SC at CP6 for the right temporal region (channels 12–15).

**a.** Spatial-specific approach For each long channel and each chromophore, the signal from the anatomically nearest SC for the same chromophore (i.e., HbO regressed with HbO, HbR with HbR) was included as a regressor of no interest in the GLM. This chromophore-specific approach follows the implementation of the nearest-SC inclusion in a GLM in Homer3^30^. The nearest SC was defined as the one selected for the same lobe in which the long channel was located.
**b.** Signal-specific approach For each long channel and each chromophore, the signal from the most correlated SC for the same chromophore (i.e., HbO regressed with HbO, HbR with HbR) was included as a regressor of no interest in the GLM. This approach follows the implementation of the most correlated SC inclusion in a GLM in Homer3^30^.
**c.** Non-selective mean-based approach Signals from the three available SCs were averaged separately for HbO and HbR. For each long channel and each chromophore, the resulting two regressors (mean SC-HbO and mean SC-HbR) were included as regressors of no interest in the GLM.
**d.** Non-selective pooled approach Raw signals from the three available SCs for both chromophores were processed using the PCA-based method described above. For each long channel and each chromophore, the resulting six components were included as regressors of no interest in the GLM.
3. Full SC availability (n = 8): In the next four analyses, the signals from all available SCs (n = 8, one per source) were used.

**a.** Spatial-specific approach For each long channel and each chromophore, the signal from the anatomically nearest SC (i.e., the one physically attached to the optode source of the long channel) for the same chromophore was used as a regressor of no interest in the GLM.
**b.** Signal-specific approach For each long channel and each chromophore, the signal from the most correlated SC for the same chromophore (i.e., HbO regressed with HbO, HbR with HbR) was included as a regressor of no interest in the GLM.
**c.** Non-selective mean-based approach Raw signals from the eight available SCs were averaged separately for HbO and HbR. For each long channel and each chromophore, the resulting two regressors (mean SC-HbO and mean SC-HbR) were included as regressors of no interest in the GLM.
**d.** Non-selective pooled approach without PCA Raw signals from the eight available SCs for both chromophores (i.e., 8 HbO and 8 HbR) were all included as regressors of no interest in the GLM. Note that no PCA was applied to the raw SC signal.
**e.** Non-selective pooled approach with PCA Raw signals from the eight available SCs for both chromophores were processed using the PCA-based method described above. For each long channel and each chromophore, the resulting sixteen components were included as regressors of no interest in the GLM.

### 2.7. Statistical analysis

We opted for a Bayesian statistical framework to analyze our data, an approach that is gaining traction for its suitability in fNIRS research^9^ and offers several key advantages for the specific goals of this methods-comparison study. First, this approach provides a more robust framework for handling the multiple comparisons inherent in our design. Specifically, Bayesian analysis offers inherent robustness to Type I error rates without requiring the power-reducing statistical corrections (e.g., Bonferroni, FDR) typical of frequentist approaches^45,46^. Second, Bayesian analysis allows for the quantification of evidence in favor of both the alternative hypothesis and the null hypothesis. This ability to find evidence for an absence of activation is essential for demonstrating when a preprocessing pipeline fails to recover an expected neural signal, a conclusion that is difficult to draw from traditional frequentist statistics^46^. By reporting Bayes Factors, we can directly assess the strength of evidence for or against our hypotheses in a way that is more intuitive and less susceptible to the pitfalls of repeated testing.

All analyses described above yielded a beta value for each condition regressor (A, V, AV), for each subject, long channel, and chromophore. These beta estimates were then submitted to Bayesian t-tests, either one-sample tests against zero or two-sample t-tests comparing conditions (see next section for details), to quantify a degree of logical support or belief for an effect of interest. We report Bayes Factors (BF₁₀) as a relative measure of evidence in favor of the alternative hypothesis compared to the null model^46^. Traditionally, a BF₁₀ between 1 and 3 is considered weak evidence for the tested model, between 3 and 10 as positive evidence, between 10 and 100 as strong evidence, and greater than 100 as decisive evidence. Similarly, to interpret the strength of evidence in favor of the null model, a BF₁₀ between 0.33 and 1 is considered weak evidence, a BF between 0.01 and 0.33 as positive evidence, a BF between 0.001 and 0.01 as strong evidence, and a BF lower than 0.001 as decisive evidence^45^.

### 2.8. Pipeline quality metrics and validation

To compare the different analysis pipelines, we computed two evaluation metrics for each:

- <u>Metric 1</u>: emergence tests; we counted the number of channels showing evidence for an effect in any condition (relative to baseline) in the expected direction. We ran emergence tests in each channel and for each condition by performing one-tailed, one-sample Bayesian t-tests against 0 across subjects. A channel was considered as showing evidence for an effect in the expected direction as follows:

- **For the temporal lobes**: BF₁₀ > 3 for condition A compared to 0 (A > 0 for HbO, A < 0 for HbR) OR BF₁₀ > 3 for condition AV compared to 0 (AV > 0 for HbO, AV < 0 for HbR)
- **For the occipital lobe**: BF₁₀ > 3 for condition V compared to 0 (V > 0 for HbO, V < 0 for HbR) OR BF₁₀ > 3 for condition AV compared to 0 (AV > 0 for HbO, AV < 0 for HbR)
- <u>Metric 2</u>: between-condition difference tests; we counted the number of channels where there was evidence for a difference between conditions in the expected direction. Comparisons between conditions were only conducted in channels that showed evidence of an effect for at least one condition relative to the baseline in the expected direction (as defined in Metric 1). Specifically, we performed pairwise comparisons between conditions in each channel selected by Metric 1, using one-tailed, two-sample Bayesian t-tests across subjects. Channels were considered showing an effect in the expected direction as follows:

- **For the temporal lobes**: BF₁₀ > 3 for condition A vs V (A > V for HbO, A < V for HbR) OR BF₁₀ > 3 for condition AV vs V (AV > V for HbO, AV < V for HbR)
- **For the occipital lobe**: BF₁₀ > 3 for condition V vs A (V > A for HbO, V < A for HbR) OR BF₁₀ > 3 for condition AV vs A (AV > A for HbO, AV < A for HbR)

These two complementary metrics were chosen to assess the performance of each analysis pipeline. Metric 1 evaluated the overall emergence of task-related activity relative to baseline, providing a global measure of the pipeline’s ability to recover expected neural responses. Metric 2, in contrast, offered a more targeted evaluation by assessing expected between-condition differences within the subset of channels already identified by Metric 1. Together, these metrics ensured that the evaluation captured both the sensitivity of the pipelines to detect expected activations relative to a no-stimulation baseline and their capacity to discriminate between conditions in anatomically relevant regions.

## 3. Results

To ensure a clear and comprehensible presentation of our findings, we will first present HbO and HbR results for all channels in the main text and in Figures 3 to 5 for three representative pipelines: the baseline ’No SCs’ condition, the ’Limited SC availability with PCA’ approach, and the ’Full SC availability with PCA’ approach. These pipelines were selected as they most effectively illustrate the progressive impact of including and optimizing SC regression. Channel-wise results and figures for the other seven pipelines can be found in the Supplementary Materials (Figures S1-S7). We will then present the quantitatively summarized conclusive performance of every tested strategy in our main validation metrics figure for HbO and HbR (Figure 6).

**Figure 3:**
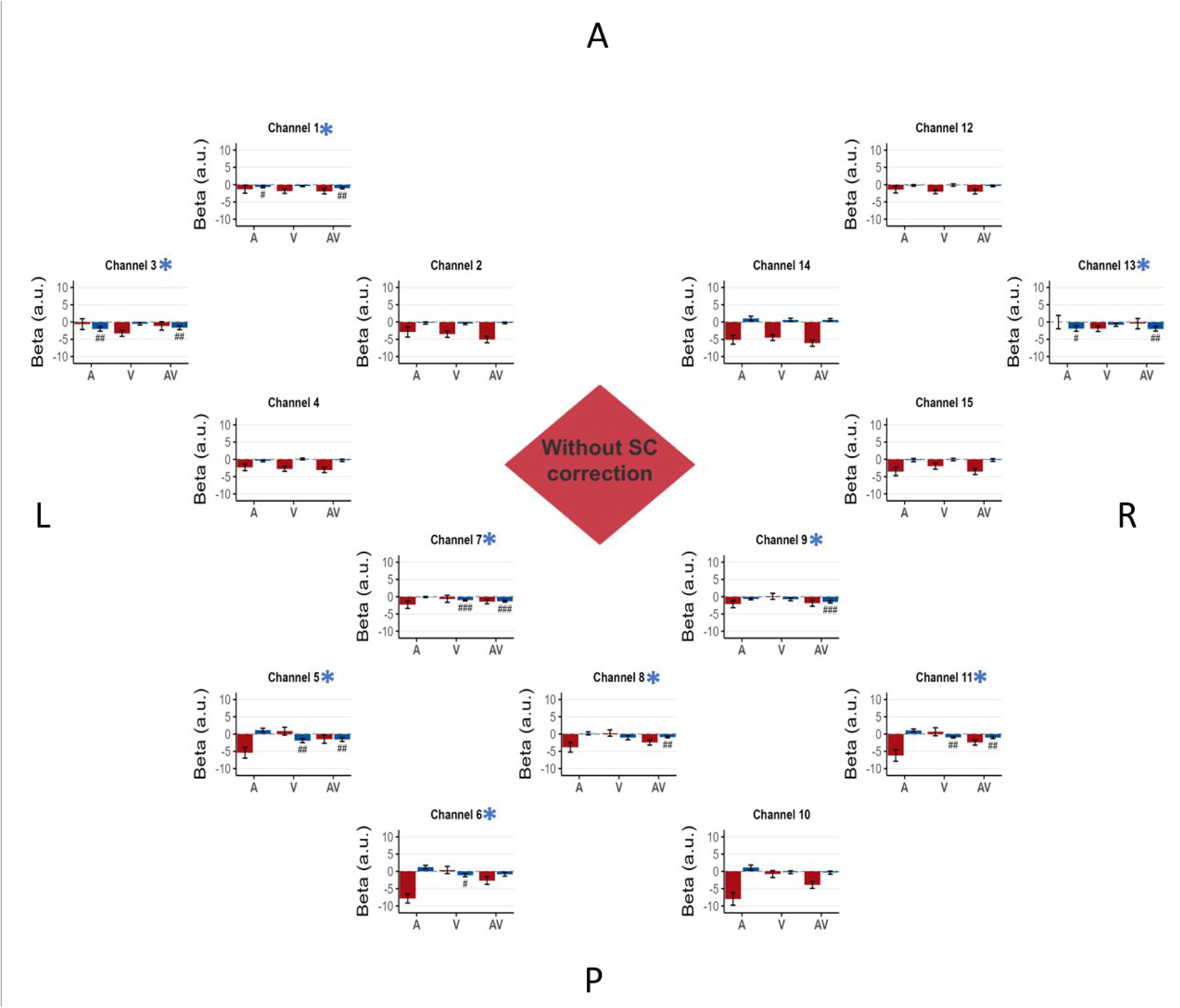
Baseline: no SC correction. In this bird’s-eye view of the fNIRS montage, channels are positioned according to their approximate location on the scalp, with labels indicating the anterior (A), posterior (P), left (L), and right (R) regions of the montage. Channel 1 to 4 are located above the left temporal auditory areas, Channel 5 to 11 are located above the occipital area, channel 12 to 15 are located above the right temporal auditory area. Each graph represents the beta (a.u.: arbitrary units) estimated from both HbO and HbR signals, averaged across the 16 participants. The bars are visually coded by chromophore and condition: the outline color distinguishes between HbO (red) and HbR (blue), while the fill color represents the stimulation type (auditory: black, visual: white, audiovisual: grey). Beta values are plotted to directly visualize the magnitude of the estimated physiological effect, while the statistical evidence is presented as follow for each channel and each condition, a # symbol is displayed when the Bayes Factor (BF₁₀) from a one-tailed, one-sample Bayesian t-test against 0 exceeds 3 in the expected direction (see method section 2.8), ## when BF₁₀ > 10, and ### when BF₁₀ > 100. Error bars indicate the standard error of the mean for each condition. A red asterisk (HbO) and/or a blue asterisk (HbR) next to the channel name indicates that the channel showed evidence of the expected effect according to Metric 1 described in section 2.8.

**Figure 4:**
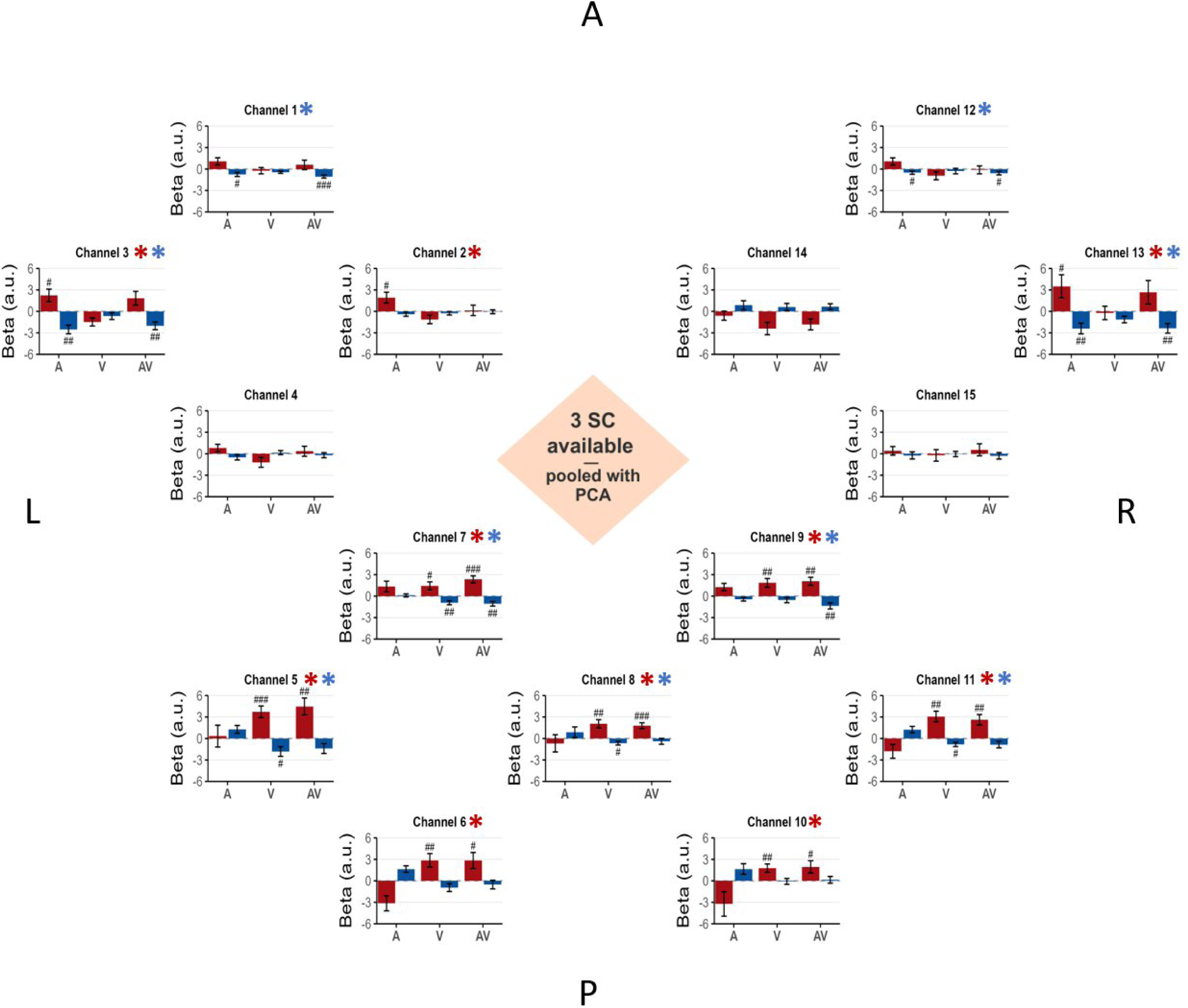
Limited short-channel availability - non-selective pooled approach. Same as in Figure 3 but showing results from the analysis using PCA-transformed components for regression in the limited short-channel availability (n = 3) scenario.

**Figure 5:**
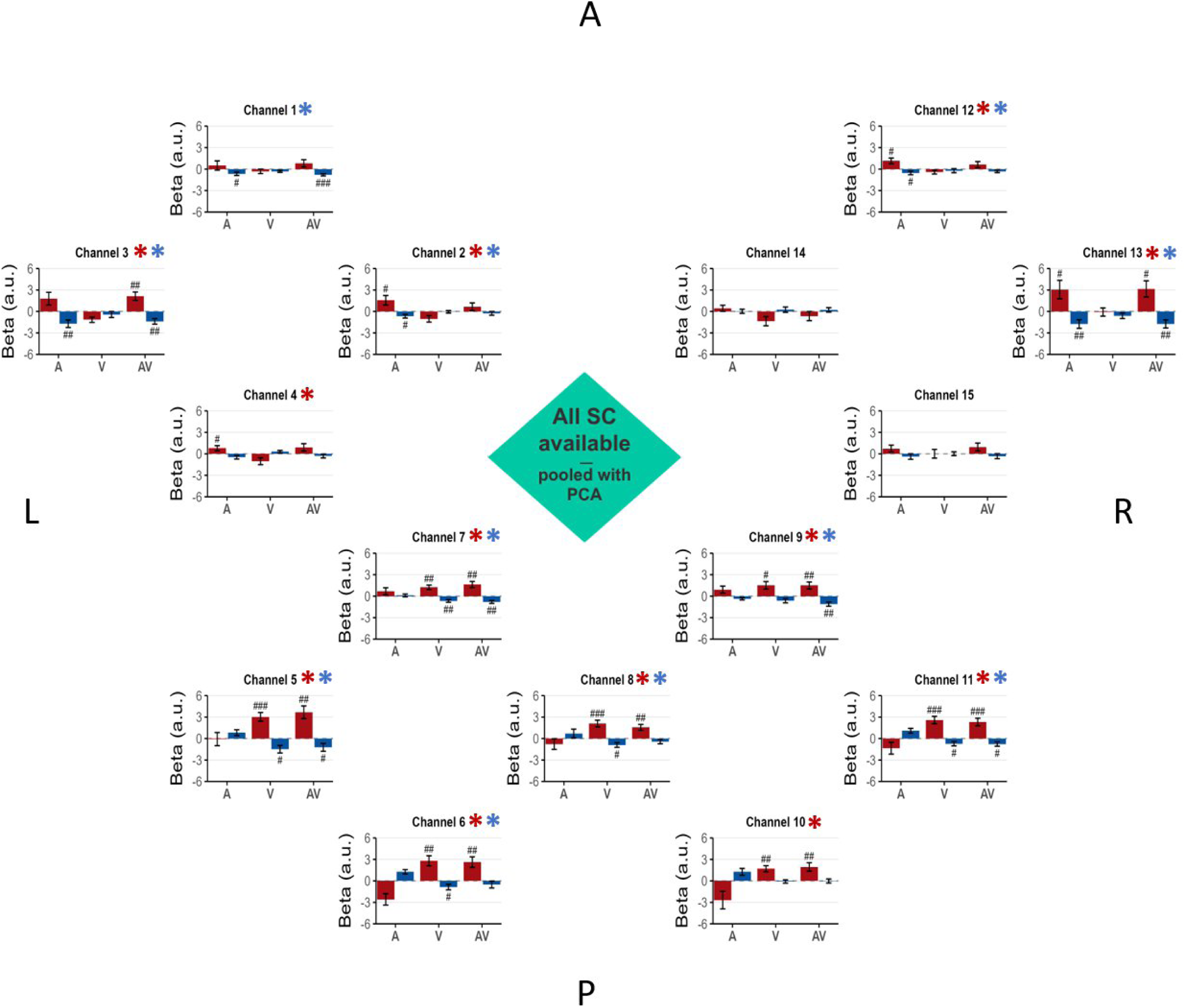
Full short-channel availability – non-selective pooled approach with PCA. Same as in Figure 4 but showing results from the analysis using PCA-transformed components for regression in the full short-channel availability (n = 8) scenario.

**Figure 6:**
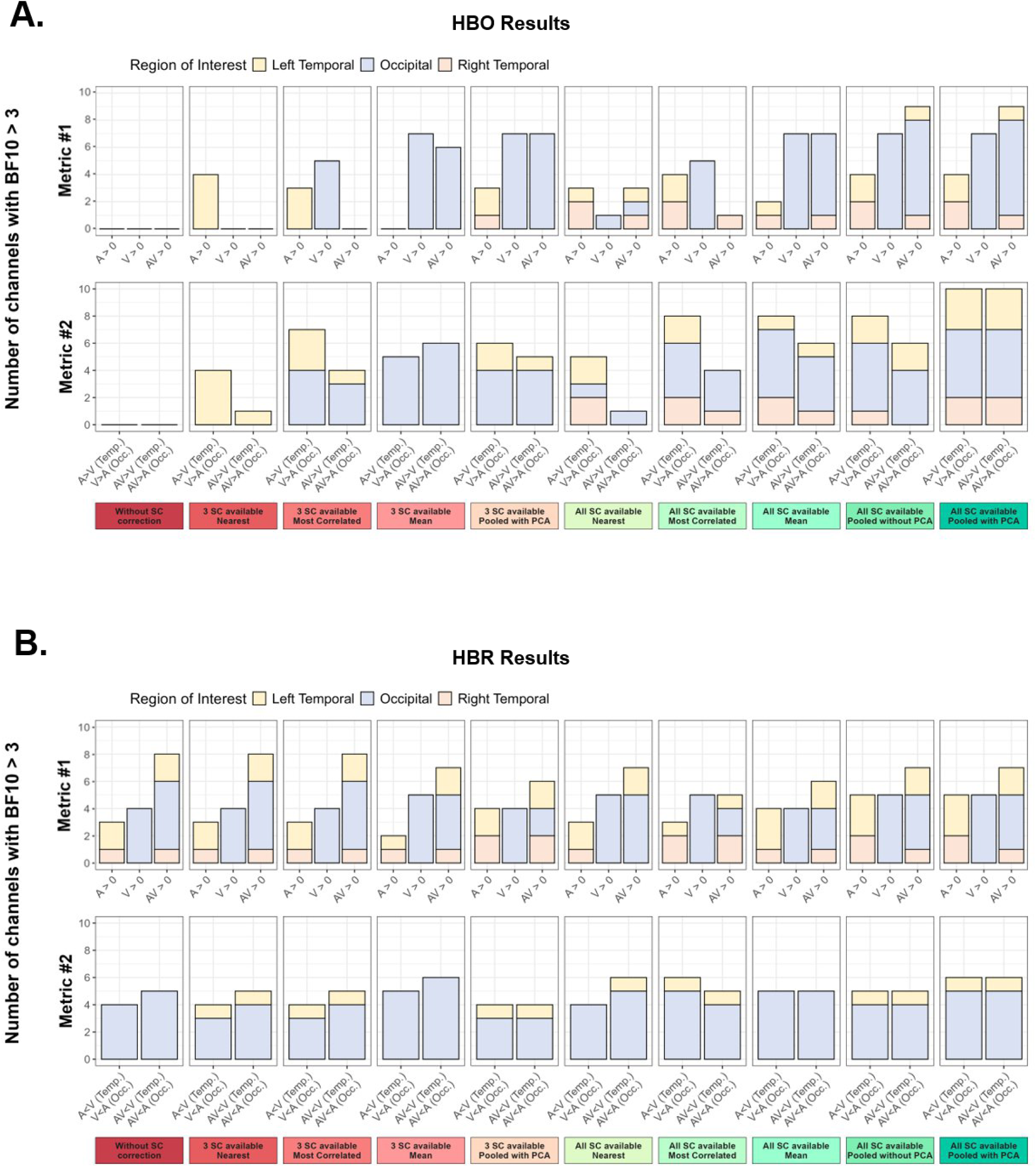
Summary of validation metrics. The figure presents the results for A) HbO and B) HbR. The top panels display, for each analysis pipeline and across all regions of interest, the number of channels showing evidence of activation (BF₁₀ > 3) based on a one-tailed, one-sample Bayesian t-test against zero (metric 1). Conditions included Auditory (A), Visual (V), and Audiovisual (AV) stimulation, and tests were directed toward positive effects (A > 0, V > 0, AV > 0). Channels were retained for further analysis only if they showed a BF₁₀ > 3 in the expected direction; specifically, A > 0 or AV > 0 in temporal areas, and V > 0 or AV > 0 in occipital areas. The bottom panels display, for each analysis pipeline and across regions of interest, the number of channels (among those showing expected emergence in metric 1) with evidence of differential activation (BF₁₀ > 3) based on one-tailed, two-sample Bayesian t-tests between conditions. The tests were directed as follows: A > V and AV > V in temporal areas, and V > A and AV > A in the occipital area.

### 3.1. Baseline: no SC correction

The beta values for the A, V, and AV conditions from the regression analysis without SC correction are presented in Figure 3. This figure displays the results for all channels for both HbO and HbR. The summary of the validation metrics is presented in Figure 6 for HbO and HbR.

For the HbO signal, most channels exhibited negative beta values in response to auditory, visual, and audiovisual stimulation. Critically, no channel in the occipital lobes or the temporal lobe showed emergence of activity in the expected direction (metric 1). Consequently, as no channel met the criteria for emergence, we did not proceed with testing for differences between conditions (metric 2).

In contrast, the HbR signal revealed detectable task-related activity even without correction. In the occipital area, four channels (5, 6, 7 and 11) showed positive evidence for emergence of activity (metric 1) in the expected direction in condition V vs. 0 (BF10 > 3), while five channels (5, 7, 8, 9 and 11) showed it for condition AV vs. 0. Among these channels, for between-condition differences (metric 2), four channels (5, 6, 7 and 11) showed strong to decisive evidence for higher beta values in condition V vs. A (31.06 < BF10 < 204.81), and five channels (5, 6, 7, 8 and 11) showed positive to decisive evidence for higher beta values in condition AV vs. A (3.5 < BF10 < 173.71). In the left temporal area, two channels (channels 1 and 3) showed evidence for emergence of activity (metric 1) in the expected direction in condition A vs. 0 and AV vs. 0 (BF10 > 3). However, those channels did not show evidence of an effect for any of the pairwise comparisons between conditions (metric 2). In the right temporal area, only channel 13 showed evidence for emergence of activity (metric 1) in the expected direction in condition A vs. 0 and AV vs. 0 (BF10 > 3), but it also did not show evidence of an effect for any of the pairwise comparisons.

### 3.2. Limited short-channel Availability (n = 3)

Our results first address the scenario simulating limited short-channel availability (n=3). In line with the rationale outlined in our methods, this section focuses specifically on the performance of the non-selective pooled approach that uses PCA. This pipeline is highlighted because it represents a global correction strategy designed to model systemic noise effectively, even under common hardware constraints. Beta values for the A, V, and AV conditions under this scenario are presented in Figure 4, which simultaneously displays the regression results for both HbO and HbR signals. The summary of the validation metrics is presented in Figure 6 for HbO and HbR.

For the HbO signal, in the occipital area, all seven channels showed evidence for the emergence of activity in the expected direction (metric 1) in conditions V vs. 0 and AV vs. 0 (BF10 > 3). Among them, for the between-condition differences (metric 2), four channels (channels 6, 8, 10 and 11) showed positive to decisive evidence for higher beta in condition V vs. A (3.03 < BF10 < 245.73), and four channels (channels 5, 6, 10 and 11) showed positive to decisive evidence for higher beta in condition AV vs. A (3.57 < BF10 < 105.89).

In the left temporal area, two channels (channels 2 and 3) showed evidence for emergence of activity in the expected direction (metric 1) in condition A vs. 0 (BF10 > 3). For the between-condition differences (metric 2), those two channels showed strong evidence for higher beta values in condition A vs. V (22.3 < BF10 < 53.4), and one (channel 3) showed strong evidence for higher beta values in condition AV vs. V (BF10 = 15.9).

In the right temporal area, one channel (channel 13) showed evidence for emergence of activity (metric 1) in condition A vs. 0 (BF10 > 3). However, this channel did not show evidence of between-condition differences (metric 2).

For the HbR signal, in the occipital area, five channels (channels 5, 7, 8 and 11) showed positive evidence for emergence of activity (metric 1) in the expected direction in conditions V vs. 0, while two channels (channels 7 and 9) showed it for condition AV vs. 0. Among them, for between-condition differences (metric 2), three channels (channels 5, 7, and 11) showed strong evidence for higher beta values in condition V vs. A (16.8 < BF10 < 96.51), and the same three channels showed strong evidence for higher beta values in condition AV vs. A (17.35 < BF10 < 32.15). In the left temporal area, two channels (channels 1 and 3) showed evidence for emergence of activity (metric 1) in the expected direction in condition A vs. 0 and AV vs. 0 (BF10 > 3). For between-condition differences (metric 2), only channel 3 showed positive evidence for higher beta values in condition A vs. V (BF10 = 5.87), and only channel 1 showed positive evidence for higher beta values in condition AV vs. V (BF10 = 4.43).

In the right temporal area, two channels (channels 12 and 13) showed evidence for emergence of activity (metric 1) in the expected direction in condition A vs. 0 and AV vs. 0 (BF10 > 3). However, those channels did not show evidence of an effect for any of the pairwise comparisons between conditions (metric 2).

### 3.3. Full short-channel availability (n = 8)

We next present detailed results under the condition of full short-channel availability (n=8). This section details the results from the non-selective pooled approach with PCA, as this pipeline leverages the complete dataset to construct the most robust data-driven model of systemic interference. Beta values for the A, V, and AV conditions under this scenario are presented in Figure 5, which simultaneously displays the regression results for both HbO and HbR signals. The summary of the validation metrics is presented in Figure 6 for HbO and HbR.

For the HbO signal, in the occipital area, all seven channels showed evidence for the emergence of activity in the expected direction (metric 1) in condition V vs. 0 and AV vs. 0 (BF10 > 3). Among them, for between-condition differences (metric 2), five channels (channels 5, 6, 8, 10 and 11) showed strong to decisive evidence for higher beta values in condition V vs. A (27.43 < BF10 < 1765.72), and the same five channels showed positive to decisive evidence for higher beta values in condition AV vs. A (6.25 < BF10 < 188.82).

In the left temporal area, two channels (channels 2 and 4) showed evidence for emergence (BF10 > 3) of activity in the expected direction (metric 1) in condition A vs. 0 and only channel 3 in condition AV vs. 0 . For between-condition differences (metric 2), those three channels showed strong evidence for higher beta values in condition A vs. V (16.88 < BF10 < 24.75), and all three showed positive to decisive evidence for higher beta values in condition AV vs. V (5.65 < BF10 < 625.71). In the right temporal area, two channels (channels 12 and 13) showed evidence for emergence (BF10 > 3) of activity in the expected direction (metric 1) in condition A vs. 0 and only channel 13 in condition AV vs. 0 . For between-condition differences (metric 2), those two channels showed positive to strong evidence for higher beta values in condition A vs. V (3.97 < BF10 < 23.65), and the same two channels showed positive evidence for higher beta values in condition AV vs. V (3.03 < BF10 < 7.04).

For the HbR signal, in the occipital area, five channels (channels 5, 6, 7, 8 and 11) showed evidence for emergence of activity (metric 1) in the expected direction in conditions V vs. 0 and four channels (channels 5, 7, 9 and 11) in conditions AV vs. 0 (BF10 > 3). Among them, for between-condition differences (metric 2), five channels (channels 5, 6, 7, 8, and 11) showed positive to decisive evidence for higher beta values in condition V vs. A (4.95 < BF10 < 286.06), and five channels (channels 5, 6, 7, 9 and 11) showed positive to decisive evidence for higher beta values in condition AV vs. A (4.19 < BF10 < 199.13).

In the left temporal area, three channels (channels 1, 2 and 3) showed evidence for emergence of activity (metric 1) in the expected direction in condition A vs. 0 and two channels (channels 1 and 3) AV vs. 0 (BF10 > 3). For between-condition differences (metric 2), only channel 2 showed positive evidence for higher beta values in condition A vs. V (BF10 = 3.05), and only channel 1 showed positive evidence for higher beta values in condition AV vs. V (BF10 = 4.74). In the right temporal area, two channels (channels 12 and 13) showed evidence for emergence (BF10 > 3) of activity (metric 1) in the expected direction in conditions A vs. 0 and only channel 13 for AV vs. 0 condition . However, those channels did not show evidence of an effect for any of the pairwise comparisons between conditions (metric 2).

### 3.4. Comparison of analysis pipelines across quality metrics

The following section presents results across all pipelines evaluated with the two quality metrics for HbO (Figure 6a) and HbR (Figure 6b).

#### HbO results

As shown in Figure 6a, the number of emerging channels detected by both validation metrics increased steadily with the introduction and the way short-channel regressors were integrated.

Without SC correction, no channel reached significance on Metric 1 (see Figure 3), and accordingly, no effect was observed on Metric 2.

When the nearest SC strategy was used under the limited SC availability scenario, a limited improvement appeared. In Metric 1, four left temporal channels emerged for the A vs 0 condition (see supplementary Figure S1) and in Metric 2, the four left temporal channels showed evidence of an effect for the A vs V and/or AV vs V comparisons. No emergence was detected in the occipital channels or in the right temporal channels.

When the most correlated SC strategy was used under the limited SC availability scenario, an improvement in the occipital channels appeared. In Metric 1, five occipital channels emerged for the V vs 0 condition and three left temporal channels emerged for the A vs 0 condition (see supplementary Figure S2). In Metric 2, four occipital channels showed evidence of an effect for the V vs A and/or AV vs A comparisons, and the three left temporal channels showed evidence of an effect for the A vs V (three channels) and/or AV vs V (one channel) comparisons.

When using the mean of SC signals under the limited SC availability scenario, a clearer step forward was observed for occipital channels but not for temporal channels. In Metric 1, the seven occipital channels emerged for the V vs 0 (seven channels) and/or AV vs 0 conditions (six channels, see supplementary Figure S3) and in Metric 2, six of these occipital channels showed evidence of an effect for the V vs A (five channels) and AV vs A (six channels) comparisons.

When using the PCA components of all SC signals under the limited SC availability scenario, an additional improvement was observed, as temporal channels now showed evidence of an effect alongside the occipital ones. In Metric 1 the seven occipital channels emerged for the V vs 0 and AV vs 0 conditions and three temporal channels (two left, one right) emerged for the A vs 0 condition (see Figure 4). In Metric 2 five occipital channels showed evidence of an effect for the V vs A and/or AV vs A comparisons, and two left temporal channels showed evidence of an effect for the A vs V (two channels) and/or AV vs V (one channel) comparisons.

When the nearest SC strategy was used under the full SC availability scenario, the overall quality of the results decreased, especially for the occipital channels. In Metric 1 only one occipital channel emerged for the V vs 0 and AV vs 0 conditions and four temporal channels (two left, two right) emerged for the A vs 0 condition (see supplementary Figure S4). In Metric 2, one occipital channel showed evidence of an effect for the V vs A and AV vs A comparisons and four temporal channels showed evidence of an effect for the A vs V (two left, two right) comparison.

When the most correlated SC strategy was used under the full SC availability scenario, the overall quality of the results slightly increased compared to the nearest approach in full SC availability, but decreased compared to the pooled PCA approach with limited SC availability. In Metric 1, five occipital channels emerged for the V vs 0 condition and four temporal channels (two left, two right) emerged for the A vs 0 condition and one right temporal channel for the AV vs 0 condition (see supplementary Figure S5). In Metric 2, four occipital channels showed evidence of an effect for the V vs A and/or AV vs A comparisons, and the four temporal channels showed evidence of an effect for the A vs V (four channels) and/or AV vs V (one channel) comparisons.

When using the mean of SC signals under the full SC availability scenario, results drastically improved. In Metric 1, for occipital channels all of them emerged for the V vs 0 condition and six of them for the AV vs 0 condition. Two temporal channels (one left, one right) emerged for the A vs 0 condition and only one temporal channel (on the right) for AV vs 0 condition (see supplementary Figure S6). In Metric 2 five occipital channels showed evidence of an effect for the V vs A comparison and/or AV vs A comparison. Three temporal channels showed evidence of an effect for the A vs V (one left, two right) and/or AV vs V (one left, one right) comparisons.

Using all raw SC signals under the full SC availability scenario produced drastically better results. In Metric 1, the seven occipital channels emerged for the V vs 0 and AV vs 0 conditions and four temporal channels emerged for the A vs 0 (two lefts, two rights) and/or AV vs 0 (one left, two rights) conditions (see supplementary Figure S7). In Metric 2, five occipital channels showed evidence of an effect for the V vs A and/or AV vs A (four channels) comparisons and three temporal channels showed evidence of an effect for the A vs V (two left, one right) and/or AV vs V (two left) comparisons.

Finally, using the PCA components of SC signals under the full SC availability scenario produced the highest-quality results. In Metric 1, results were the same as the previous analysis (see Figure 5). In Metric 2, five occipital channels showed evidence of an effect for the V vs A and AV vs A comparisons and five temporal channels showed evidence of an effect for the A vs V and AV vs V comparisons (three left, two right).

#### HbR results

As shown in Figure 6b, HbR signal was less affected by the introduction and the way short-channel regressors were integrated.

In Metric 1, a minimum of eight channels consistently emerged in the expected direction across all analysis pipelines. The *Limited short-channel availability – spatial-specific approach* (Figure S1) yielded the fewest emerging channels, with five occipital and three temporal (two left, one right), whereas the *Full short-channel availability – non-selective approach with PCA* (Figure 5) yielded the most, with six occipital channels and five temporal channels (three left, two right) showing effects in the expected direction.

In Metric 2, a minimum of five channels consistently showed evidence of an effect for the between-condition comparisons across all analysis pipelines. The *Baseline: no SC correction* approach and the *Limited short-channel availability – non-selective* approach yielded the fewest between-condition effects, with the former showing five occipital channels and no temporal channels, and the latter showing an effect in three occipital and two left-temporal channels. In contrast, the *Full short-channel availability – non-selective approach with PCA* yielded the most, with six occipital channels and two left-temporal channels showing an effect.

Overall, similar to the HbO signal, HbR results showed that in both metrics the non-selective approach using all PCA components of the raw short-channel signals yielded the most consistent outcomes. However, unlike HbO, the absence of SC regression had less drastic effects, with some expected effects—albeit weaker—still observable even without SC correction.

## 4. Discussion

The goal of the present study was to systematically assess the impact of SC regression on statistical outcomes in fNIRS studies, using an ecological audiovisual paradigm designed to elicit distinct activation patterns in the temporal and occipital cortices. By comparing ten GLM-based preprocessing strategies that varied in the number, anatomical specificity, and dimensionality reduction of SC signals, we aimed to identify the most effective pipeline for isolating neural activity from extracerebral physiological noise captured with SCs. To robustly evaluate each strategy’s efficiency, we relied on two complementary Bayesian metrics. The first metric assessed the emergence of task-related activity in the expected direction, allowing us to identify channels that exhibited emergence of activation consistent with the experimental design. The second metric, applied only to the subset of channels identified via the first, quantified the strength and reliability of between-condition differences. With the HbO signal, our results revealed that the absence of SC regression led to widespread negative beta estimates and no detectable emergence of neural activity in expected regions, highlighting the critical impact of physiological noise on the fNIRS signal in this context. In contrast, most pipelines that included SC regression produced zero-centered, interpretable beta distributions, enabling detection of the expected task-related activity. When comparing a spatially specific approach (using the nearest SC) or a signal-specific approach (using the most correlated SC) to a non-selective approach (averaging the raw SC signal or pooling all SCs together) in the GLM, we found that the non-selective method yielded more robust performance across both Bayesian metrics, regardless of whether a limited number of SCs or all available SCs were used. Finally, under conditions of full SC availability, applying a dimensionality reduction technique to the SC signals yielded a slight additional improvement in results, suggesting that identifying shared components across SCs, rather than leaving them mingled in variable proportions, leads to a more accurate estimation of extra-cerebral signals and enhance denoising without compromising spatial specificity. Importantly, while the HbR signal was overall less impacted by the presence or absence of SC correction, similar trends were observed, and the same SC regression strategy that maximized HbO signal interpretability also yielded the most robust and consistent results for HbR.

In line with best practice recommendations^47^, we computed block-averaged HbO and HbR signals, averaged across ROIs, and displayed them alongside their corresponding GLM-based results for temporal (Figure 7) and occipital (Figure 8) channels. Descriptively, the block-averaged signals closely matched the GLM outcomes for both chromophores: the higher the GLM betas, the more the averaged curves resembled a canonical HRF, with positive curves corresponding to positive betas and negative curves to negative betas. This was particularly evident for HbO, where introducing SC regressors yielded the most positive outcomes: the more efficient the pipeline, the more the block-averaged responses took the shape of positive, HRF-like curves in the expected directions, both in the occipital and temporal channels. The block-averaged signals thus closely matched the GLM outcomes, supporting the validity of our approach for both HbO and HbR.

**Figure 7:**
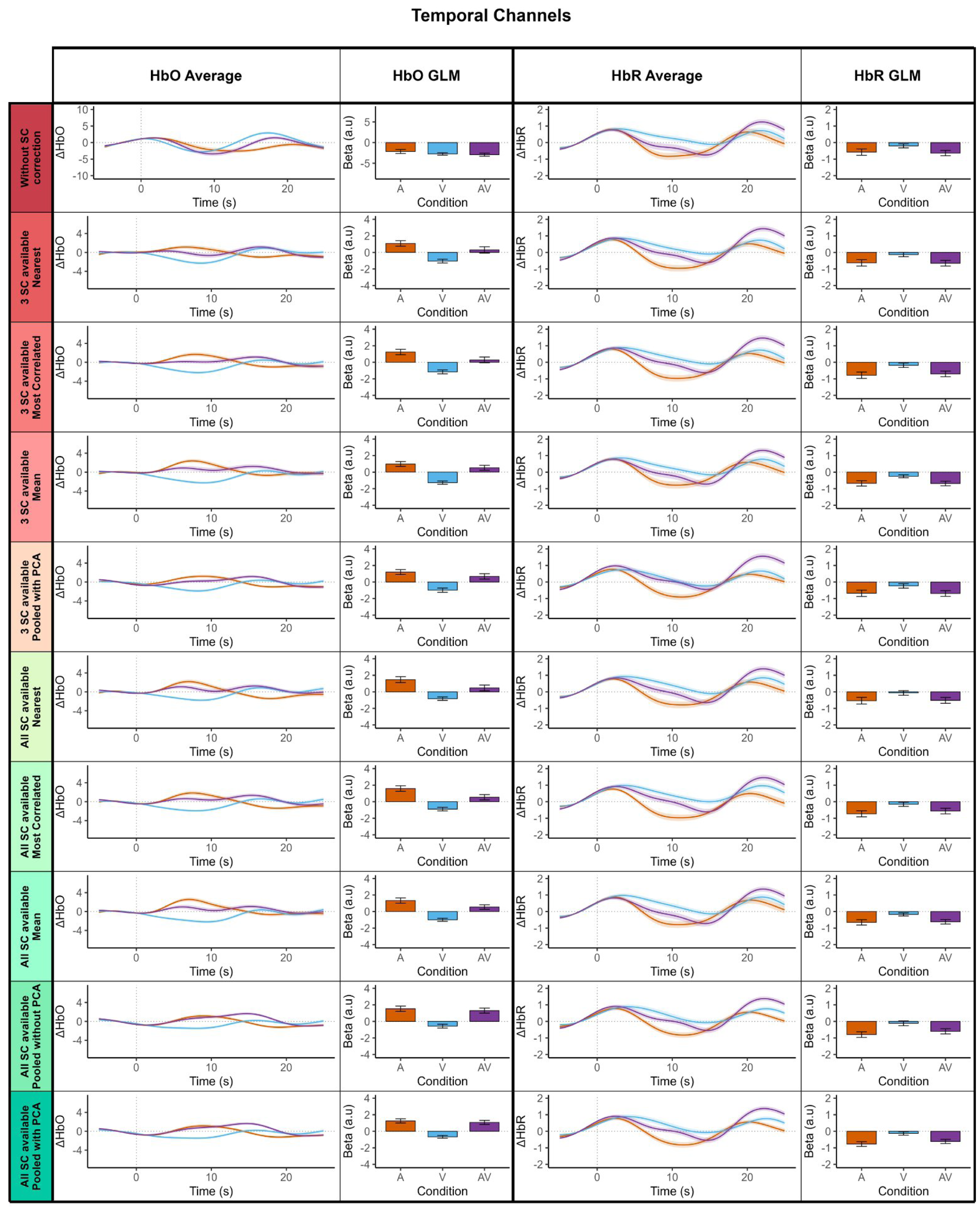
Comparison of mean hemodynamic responses and GLM beta estimates across all SC regression pipelines for the HbO and HbR signal in temporal channels. Each row displays results for one of the ten tested short-channel (SC) regression strategies, for the temporal channels (combining left and right temporal areas, channels 1-4 and 12-15). Left columns display for HbO and right columns for HbR. For each chromophore, the ’Average’ panel shows the mean hemodynamic time course calculated using the Brain AnalyzIR toolbox in MATLAB together with the Homer2 function hmrBlockAvg^30^. As in the GLM-based analyses, the same preprocessing and denoising steps were applied, including short-channel regression with identical parameters. Short-channel correction was performed prior to averaging, so that the resulting time courses reflected the corrected signals. A time window from –5 to 25 s relative to stimulus onset was used and baseline correction was applied by mean-centering the signal over the –5 to 0 s interval. The ’GLM’ panel shows the corresponding mean beta estimates. For both approaches, results are averaged across all 16 participants and all channels within the region. Stimulation conditions are color-coded: Auditory (A) in orange, Visual (V) in blue, and Audiovisual (AV) in purple. Standard errors are represented as colored ribbons around their corresponding curves for the Average approach, and as error bars for the GLM approach.

**Figure 8:**
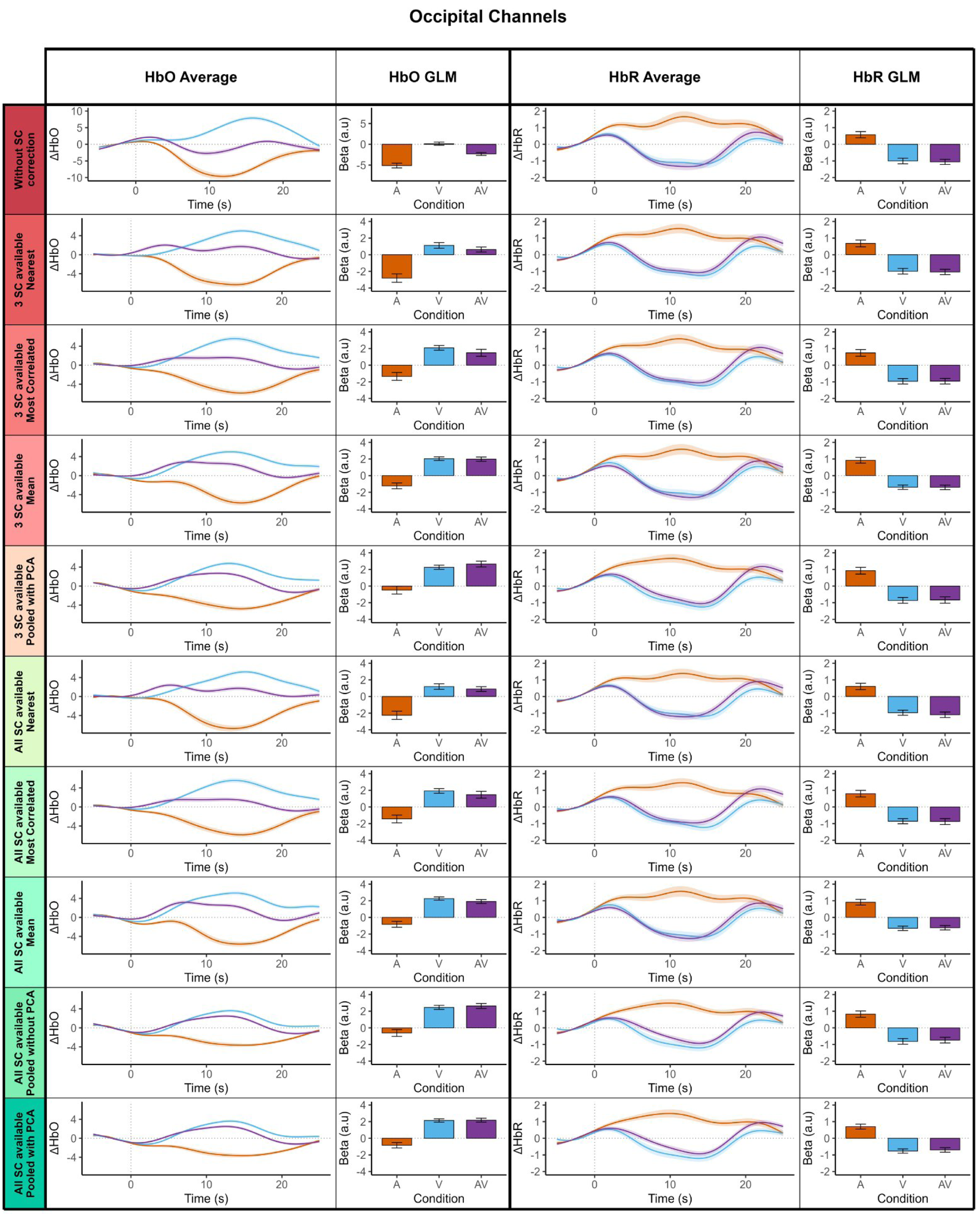
Comparison of mean hemodynamic responses and GLM beta estimates across all SC regression pipelines for the HbO and HbR signal in occipital channels. Same as Figure 7 but for occipital channels (channels 5-11).

### 4.1. The impact of omitting SC regression

In the specific context of our block-design, audio-visual task in healthy adults, our results indicate that omitting SC regression in GLM-based analysis can markedly affect the interpretability and reliability of fNIRS results. When no SC correction was applied to the HbO signal, beta estimates were systematically negative across the cortex, with no detectable emergence of neural activity in regions where activation was expected based on the task design (Figure 3). None of the channels passed the criteria of our first metric (i.e., emergence in the expected direction) when the SC signal was not included in the regression (Figure 6). These findings suggest that, in this experimental context, the SC-uncorrected pipeline failed to recover the expected neural activation from the data.

This failure of the uncorrected pipeline is consistent with previous studies showing that signal cleaning, particularly using SC regression, is essential for recovering task-related neural activity. SC regression consistently outperforms uncorrected pipelines across a variety of paradigms. Whether based on averaging, individual SC modeling, or principal component decomposition of SC signals, regressing out superficial components using SC reliably improves signal quality and interpretability^23,25,27^. It also surpasses alternative correction methods such as common average referencing or global channel regression, in the case of motor tasks targeting sensorimotor areas^24^. When SCs are not available, applying any regression method is still markedly better than leaving the signal uncorrected. Global signal regression^28^, ICA-based modeling of global scalp hemodynamics^20^, or spatial filtering technique^48^ all result in improved signal quality compared to raw data^26^. Although our study did not directly compare SC regression with other correction methods, our results, combined with extensive prior evidence, support the conclusion that SC regression remains a robust and reliable approach to isolating neural activity in fNIRS^23–25,27,28,47^.

### 4.2. Selective vs. non-selective SC regression under limited and full SC availability

We compared two GLM-based approaches to SC regression: two selective methods, which either assigns the nearest SC (spatial-specific) or the most correlated SC (signal-specific) to each long channel, and two non–selective methods. The first non–selective method averages the raw signals of all SCs separately for HbO and HbR and regresses both onto each long channel, while the second regresses out from each long channel all components of a principal component analysis (PCA) applied to the signals of all SCs, with both chromophores combined. While the selective approaches assume that local SCs or signal-similar SCs best reflect physiological noise affecting long channels, the non-selective methods assume that common noise components can be estimated more globally across SCs.

In both the selective methods, we regressed the long channel signal with the signal from the nearest SC or the most correlated SC in a chromophore-specific manner (i.e., HbO signals were regressed using the nearest HbO SC, and similarly for HbR) following the implementation used in Homer 3^30^. Notably, most published studies applying SC regression do not specify whether their regressors are chromophore-specific, and reporting this methodological detail would be a valuable practice to adopt in future work.

Under limited SC availability (n = 3; one SC per lobe), the spatial-specific method recovered auditory-related responses only in the left temporal region, with no channel showing emergence in right temporal or occipital areas. The signal-specific method replicated left temporal effects (though slightly weaker) and was able to recover a few visually-related responses in the occipital region. In contrast, the non-selective method using the mean of raw SC signals recovered only visually-related responses in the occipital region while no auditory-related responses were found in the temporal regions. The non-selective method using PCA components, however, not only replicated left temporal effects (though slightly weaker) but also showed emerging activity in the right temporal lobe and robust responses across the occipital cortex, with four out of seven channels meeting strong to decisive evidence thresholds for between-condition effects (V > A, AV > A). These results indicate that even with minimal SC availability, pooling SC signals and using their shared variance across channels can more effectively capture and remove systemic noise than relying on local proximity, signal similarity, or averaged raw SC signals. Using all available SCs also compensates for poor signal quality of one or multiple SCs, for example due to suboptimal scalp contact or hair interference, and reduces the risk of errors caused by relying on a single, potentially noisy SC.

Under full SC coverage (n = 8), the non-selective PCA-based approach led to the clearest and most widespread emergence of task-related responses across all regions. It revealed strong between-condition effects in the left and right temporal cortices as well as in the occipital lobe, showing broad spatial coverage and consistent activation patterns. Notably, the spatial-specific method under full SC availability failed to reach the performance of the non-selective PCA-based method under full SC availability and even fell short of the non-selective PCA-based method under limited SC availability. The signal-specific method under full SC availability showed intermediate performance: while it fell short compared to all non-selective approaches under full SC availability, it was able to recover more consistent effects across both metrics than the non-selective approach under limited SC availability. The non–selective mean-based method produced results more similar to the PCA-based approach, although it was less effective at capturing auditory-related responses in the temporal regions. These findings suggest that spatial proximity or signal similarity of SCs may not fully capture distributed physiological noise, and that aggregating shared variance, either by averaging SC signals or using PCA components across all SCs, provides a more robust correction, with the PCA-based approach being particularly effective at recovering auditory-related responses in the temporal regions. To our knowledge, the only study that directly compared spatial-specific and non-selective approaches in the sensorimotor cortex during a hand-grasping task reported comparable GLM performance between the two methods^25^. However, their non-selective analysis relied on a single PCA component, which likely limited its effectiveness, as previous work has shown that including mean SC signals or multiple PCA components is critical for capturing a broader range of systemic noise^23,27^.

When comparing limited vs. full SC availability, the non-selective PCA-based approach recovered qualitatively similar activation patterns in the occipital cortex across both conditions, indicating that visual-related effects can be detected even with only three SCs. However, the temporal lobes, particularly the right hemisphere, showed a marked improvement with full SC coverage: more channels passed our statistical criterion, and auditory-driven responses were stronger and more spatially extensive. This observation suggests that while limited SC availability can be sufficient to recover robust, widespread effects, full coverage becomes essential when targeting weaker or more spatially focal responses (here the auditory activations), in line with Klein et al^24^ who showed that including all SCs in a non-selective GLM yielded better signal quality than using just two SCs (one per hemisphere) during a motor task.

Taken together, our results indicate that non–selective methods consistently outperform selective approaches, both under limited and full SC coverage, and that the PCA-based approach produces more consistent results than averaging the raw SC signals. While the mean-based approach, implemented by default in the MNE-NIRS library^23^, and available as an option in Homer3^30^, showed comparable performance to PCA-based regression in a previous study using passive auditory tasks, this was not the case in the present study for channels in the temporal region. For the occipital region, however, both approaches yielded similar results. This difference likely reflects the fact that Luke et al.^23^ had twelve temporal channels capable of capturing auditory-related activity, whereas we had only eight bilaterally. A possible explanation is that, with limited temporal-lobe coverage, the PCA-based approach is better able to capture subtle, distributed fluctuations in the SC signals that contribute to auditory-related responses.

These findings support existing recommendations to incorporate multiple sources of systemic signal when applying SC correction^23,25,27^, and suggest that global physiological noise is more effectively addressed by modelling shared variance across multiple SCs rather than relying solely on spatial proximity or signal similarity.

Importantly, our results also highlight that the relative performance of each pipeline is not uniform across regions or tasks. For example, when SC availability was limited, the spatially specific nearest-SC approach mainly recovered left temporal auditory responses, the non-selective mean-based approach mostly captured occipital visual responses, and the signal-specific approach recovered some responses in both occipital and temporal regions. Only the PCA-based non-selective method successfully recovered both occipital and temporal activations, demonstrating that pipeline efficacy can depend on the combination of task, region, and SC availability. Under full SC coverage, non-selective PCA-based regression consistently yielded the most widespread and robust effects across both occipital and temporal regions, but the spatial-specific approach still performed relatively better for temporal channels than occipital ones, suggesting that certain tasks or regions (like temporal locations for auditory stimulation in the present case) may benefit more from local SC regression. These observations emphasize that while PCA-based non-selective regression provides the most generalizable correction, specific pipelines may still be advantageous in particular contexts, highlighting the importance of considering both the targeted brain region and task characteristics when selecting a preprocessing strategy.

### 4.3. PCA in non-selective SC regression

To our knowledge, this study is the first to directly address whether applying PCA to SC signals improves the effectiveness of non-selective GLM-based denoising. While recent work has widely adopted PCA-transformed SC regressors to account for systemic noise^23–28,49^, no prior study has explicitly compared PCA-based and raw-signal approaches using the same dataset and analysis pipeline. Our results show that PCA-based regression outperforms the use of raw SC signals, in terms of the number of channels showing condition-specific effects and the statistical strength of those effects. The raw-signal approach did recover expected activity, especially in the occipital region, but in fewer channels and with reduced evidence of strength in the temporal lobes. These results suggest that directly entering all SC signals into the GLM may introduce redundancy and multicollinearity, limiting sensitivity. Crucially, PCA offers an orthogonal representation of the shared systemic signal variance across SCs, enhancing the model’s ability to isolate task-related neural signals. Our findings support the use of PCA as the preferred method for non-selective SC regression when full SC coverage is available and robust denoising is desired.

### 4.4. Correction strategies for HbR signal

The HbR signal, analyzed using the same procedures as for HbO (except for directionality of Bayesian tests), showed consistent patterns of negative task-related responses across SC correction scenarios. Without SC correction, occipital channels showed the strongest effects, with six channels exhibiting emergence (metric 1), and up to five channels showing strong to decisive evidence for V > A or AV > A (metric 2). With limited SC availability (n=3), selective and non-selective approaches recovered similar emergence patterns, additionally uncovering between-condition differences for one channel over the left temporal area for the non-selective approaches. Full SC availability (n=8) revealed similar emergence detection but better between-condition differences detection with up to five channels in the occipital cortex for V < A and AV < A comparisons and one channel in the left temporal area for both A < V and AV < V comparisons. Overall, as with HbO, the analysis pipeline that yielded the best results was the one including all principal components of all SCs in the case of full SC availability; however, the advantage is markedly less pronounced than for HbO, as the expected effects can still be retrieved even without SC regression. Our findings are consistent with a substantial body of research showing that HbR is inherently less affected by systemic physiological noise than HbO^14,15,27,50,51^. HbR exhibits more linear and predictable response characteristics^52^, likely due to the greater sympathetic innervation of arterioles compared to venules^15^ and its lower sensitivity to corpuscular volume changes in the vasculature^52^. As a result, SC regression techniques tend to produce more pronounced improvements for HbO, while HbR typically shows less need for such correction^27,51^. Nonetheless, our results indicate that SC regression still provides a modest but consistent improvement for HbR, and that the pipeline yielding the best results for HbO likewise offers the most robust outcomes for HbR. These findings support the recommendation that, despite its lower sensitivity to systemic noise, HbR should be corrected using the same SC regression strategy as HbO to ensure optimal data quality and interpretability.

### 4.5. Limitations

Several limitations of the present study should be acknowledged. First, the specific task, stimulation paradigm, and optode montage may limit the generalizability of our findings to other cortical regions, experimental designs, or participant populations. Future work should assess whether the observed effects of different SC regression strategies hold across diverse fNIRS setups, tasks, and groups. Second, our pipeline did not include a direct comparison with SPA-NIRS procedures, a method shown to recover important systemic physiological information that cannot be captured by short-separation channels alone^51,53^. Future studies should evaluate the impact of SPA-fNIRS both as a replacement for SC regression and in combination with SC-based approaches to determine which configuration provides the most effective denoising of fNIRS signals. Finally, we employed an OLS-GLM rather than alternative methods such as AR-IRLS^54^, primarily for computational efficiency and because OLS remains widely used in both fMRI and fNIRS research^1^. Although AR-IRLS has been shown to outperform OLS overall^27^, there is no evidence that the choice between OLS and AR-IRLS interacts with different SC regression strategies. Given these considerations, OLS was considered a valid and suitable approach for the present study, though we acknowledge that alternative GLM implementations could have been reasonably chosen as well.

### 4.6. Recommendations and conclusion

To our knowledge, this is the first study to provide concrete recommendations for SC regression in fNIRS preprocessing using ecological passive auditory, visual, and audiovisual stimulation within the same participants. This cross-modal, multi-condition design enhances the generalizability of our findings across sensory modalities and cortical regions.

Our results confirm that for HbO signal, omitting SC regression leads to uninterpretable GLM outcomes, with no detectable task-related activation, underscoring that SC correction is crucial for recovering meaningful neural signals. When SCs are limited, pooling all available SC signals (rather than relying solely on spatially nearest SC, most correlated SC or averaged raw SC signals) improves detection of expected activations. Furthermore, when full SC coverage is available, applying dimensionality summarization techniques such as PCA to the pooled SC signals further enhances the ability to isolate task-related activity by capturing shared systemic signal variance and reducing redundancy; while a mean-based approach of the raw SC signals yields comparable results for visually related responses in the occipital region, the PCA-based method outperforms it in recovering auditory-related responses in the temporal areas. The PCA-based SC regression strategy should also be applied to the HbR signal, which, although less affected by systemic interference, still benefits from such correction.

Taken together, these findings, alongside other studies, support the recommendation to include all available SC signals in the GLM and apply orthogonalization techniques such as PCA, which, in the absence of additional physiological measurements like those provided by SPA-NIRS, offers a robust and effective approach for denoising fNIRS data.

## Code, Data, and Materials Availability

The anonymized preprocessed data and analytic code necessary to reproduce the analyses presented in this paper is publicly accessible at the following URL: https://github.com/yanndavlem/nirs_article.

## Acknowledgments

This work was supported by an ANRT (Association Nationale de la Recherche et de la Technologie) - Advanced Bionics CIFRE (Conventions industrielles de formation par la recherche) PhD funding (N°2022/0341) to YL, the AgeHear ANR funding (ANR-20-CE28-0016) to PB, the recurrent funding of the CNRS (Centre national de la recherche scientifique), France, and the DRCI of the CHU Hospital Toulouse. This work was also funded by a “Pack Ambition Recherche” (COGAUDYS project) from the Auvergne-Rhône-Alpes Region, awarded to AC.

ChatGPT-4 was used for purposes of language and grammar clean-up exclusively

We thank all of our subjects for their participation in the study, and the ENT team at Purpan hospital, Toulouse, for their help. The authors would also like to acknowledge Laure Arnold and the AB team for their support.

