## Supplemental Material for "Optimizing short-channel regression in fNIRS: an empirical evaluation with ecological audiovisual stimuli"

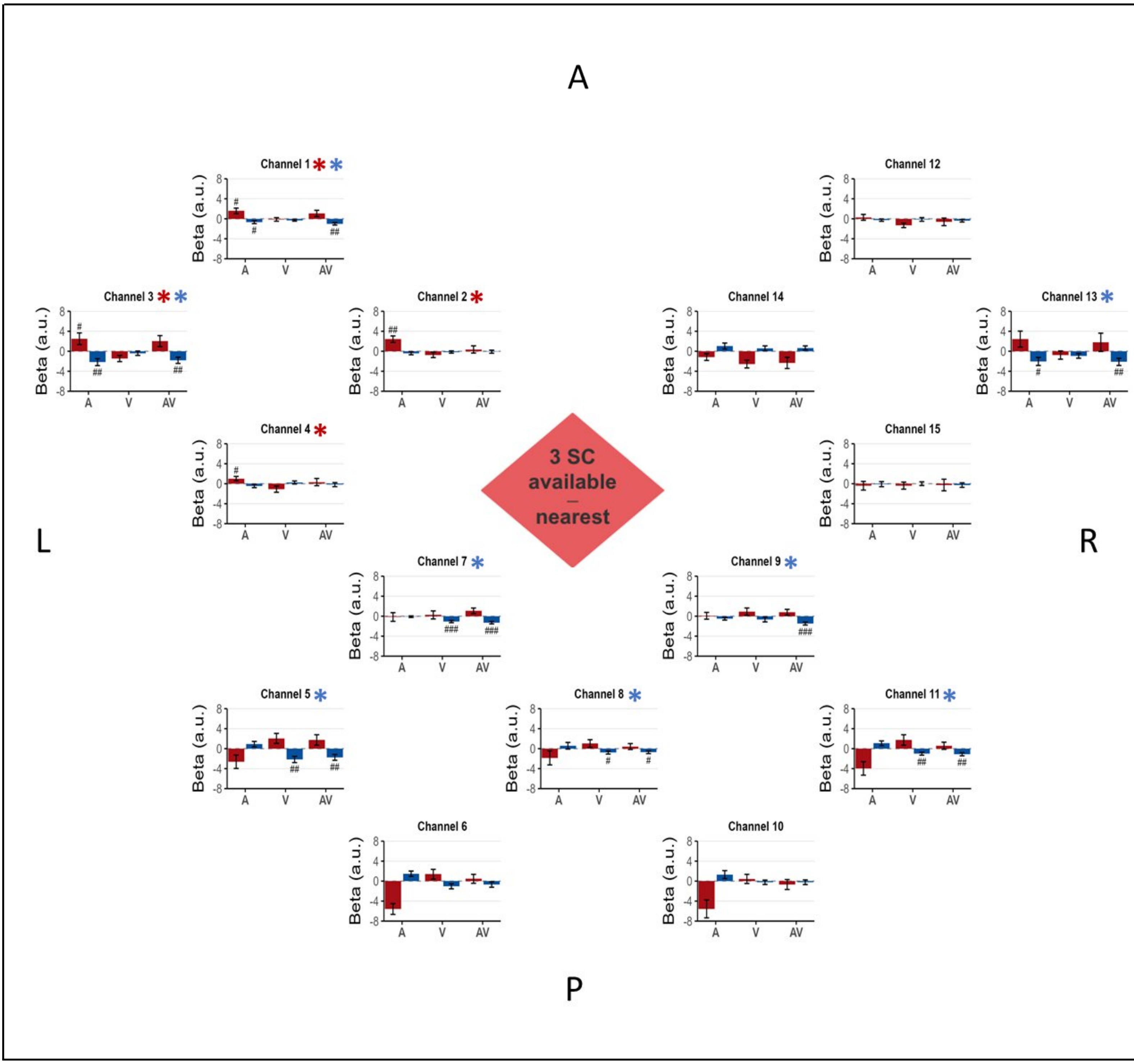


**<u>Figure S1:</u> Limited short-channel availability - spatial-specific approach (nearest).** *In this bird's-eye view of the fNIRS montage, channels are positioned according to their approximate location on the scalp, with labels indicating the anterior (A), posterior (P), left (L), and right (R) regions of the montage. Channel 1 to 4 are located above the left temporal auditory areas, Channel 5 to 11 are located above the occipital area, channel 12 to 15 are located above the right temporal auditory area. Each graph represents the beta estimated from both HbO and HbR signals, averaged*

*across the 16 participants. The bars are color-coded by chromophore, distinguishing between HbO (red) and HbR (blue). Conditions are arranged from left to right: Auditory, Visual, and Audiovisual. Beta values (a.u.: arbitrary units) are plotted to directly visualize the magnitude of the estimated physiological effect, while the statistical evidence is presented as follow for each channel and each condition, a # symbol is displayed when the Bayes Factor ($BF_{10}$) from a one-tailed, one-sample Bayesian t-test against 0 exceeds 3 in the expected direction (see method section 2.8), ## when $BF_{10}$ > 10, and ### when $BF_{10}$ > 100. Error bars indicate the standard error of the mean for each condition. A red asterisk (HbO) and/or a blue asterisk (HbR) next to the channel name indicates that the channel showed evidence of the expected effect according to Metric 1 described in Section 2.8.*

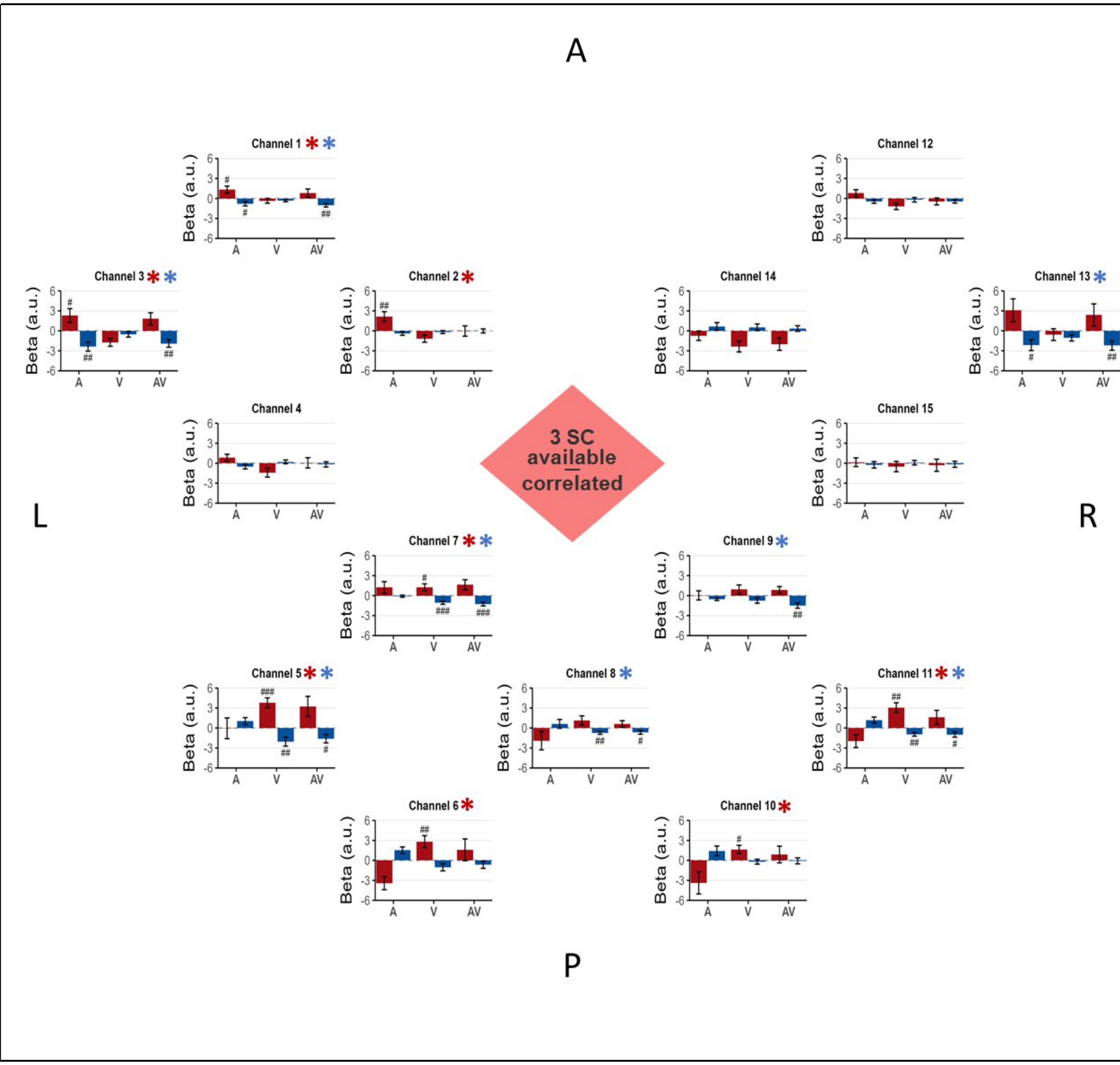


**<u>Figure S2:</u> Limited short-channel availability – signal-specific approach (most correlated)**. *Same as in Figure S1, but showing results from the analysis using the most correlated SC available for regression in the limited short-channel availability (n = 3) scenario.*

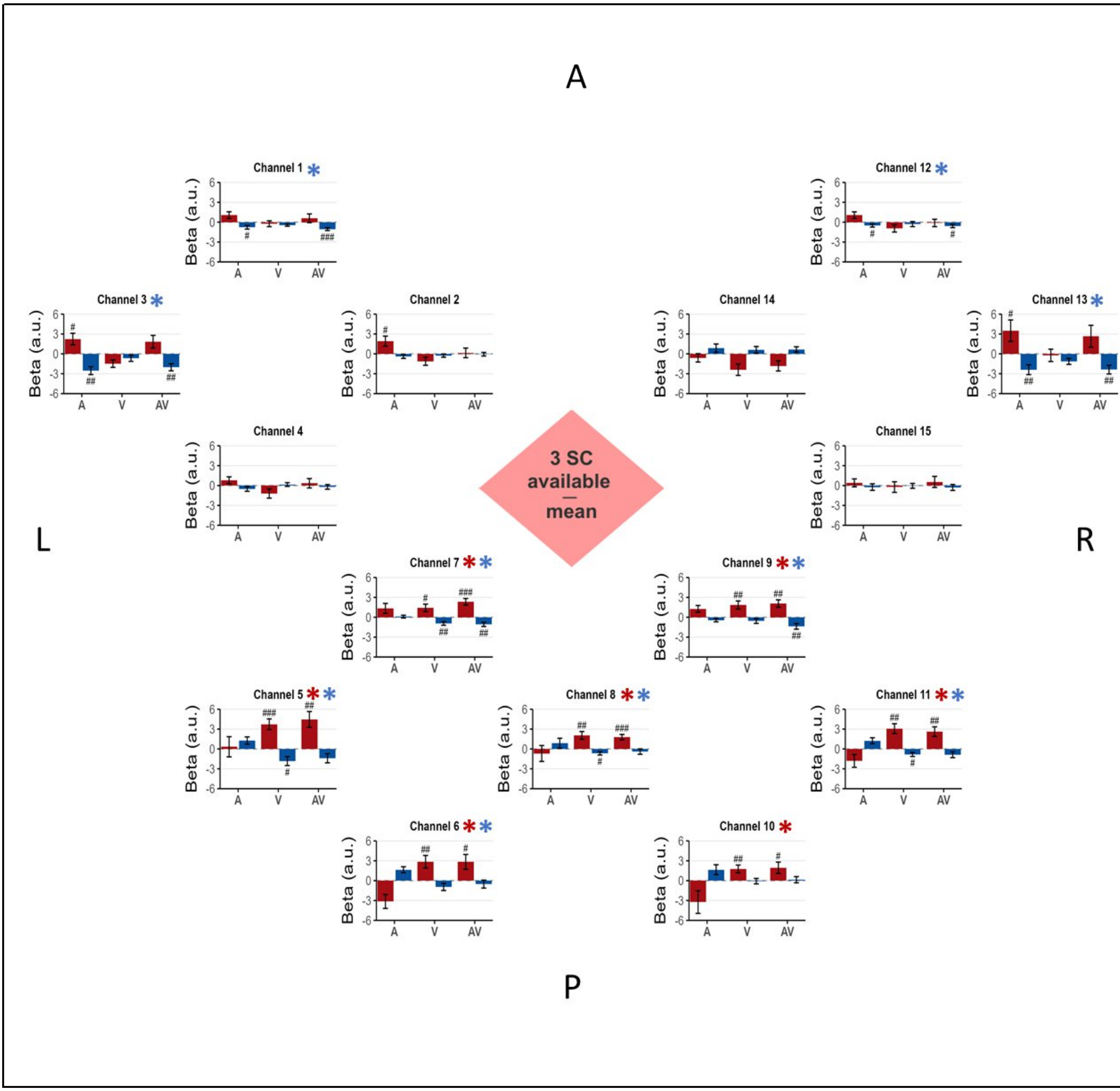


**<u>Figure S3:</u> Full short-channel availability - non-selective mean-based approach**. *Same as in Figure S1, but showing results from the analysis using the mean of all SCs available for regression in the limited short-channel availability scenario.*

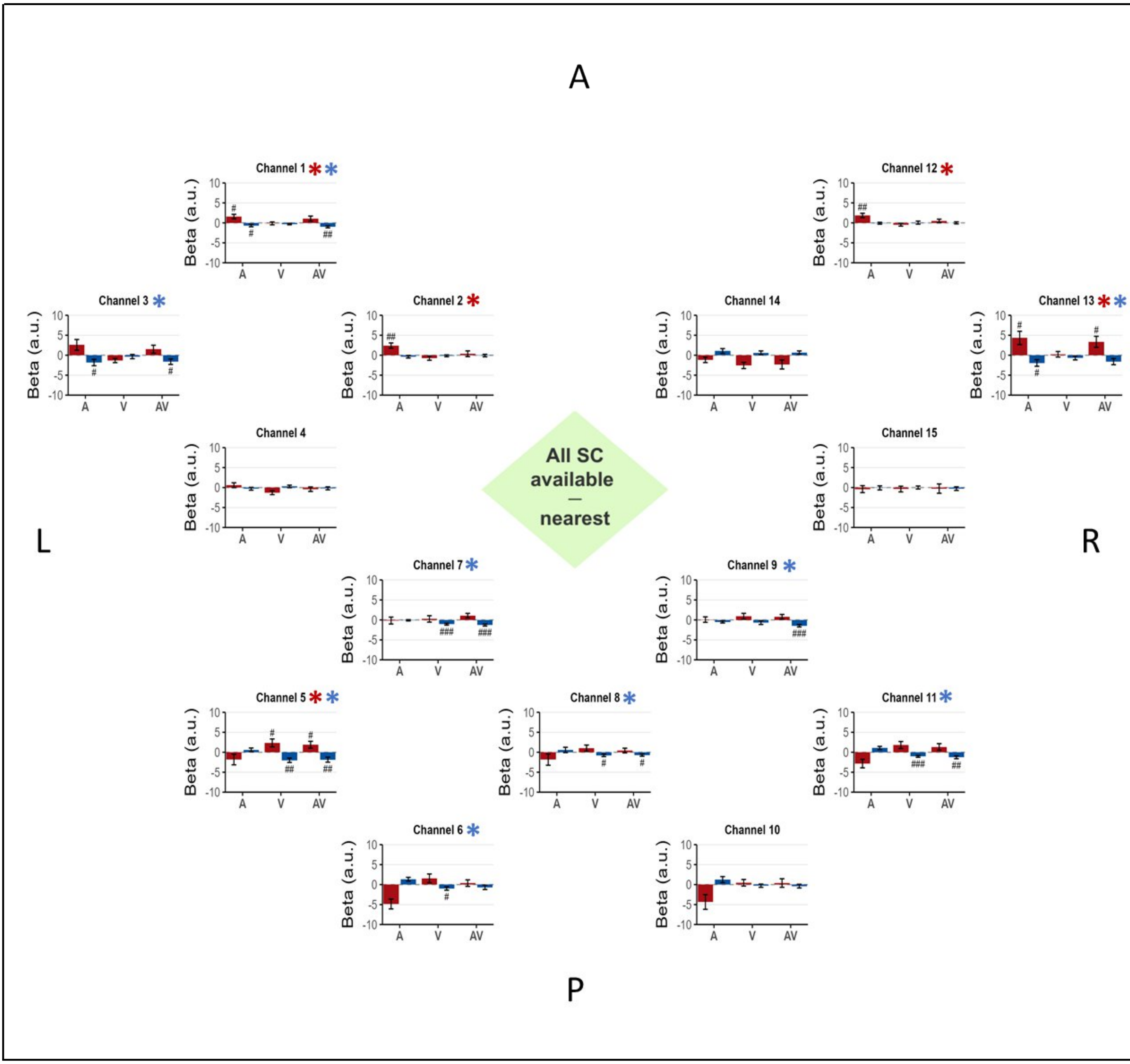


**<u>Figure S4:</u> Full short-channel availability - spatial-specific approach (nearest)**. *Same as in Figure S1, but for the scenario with full short-channel availability and using the nearest short-channel to each long-channel for regression.*

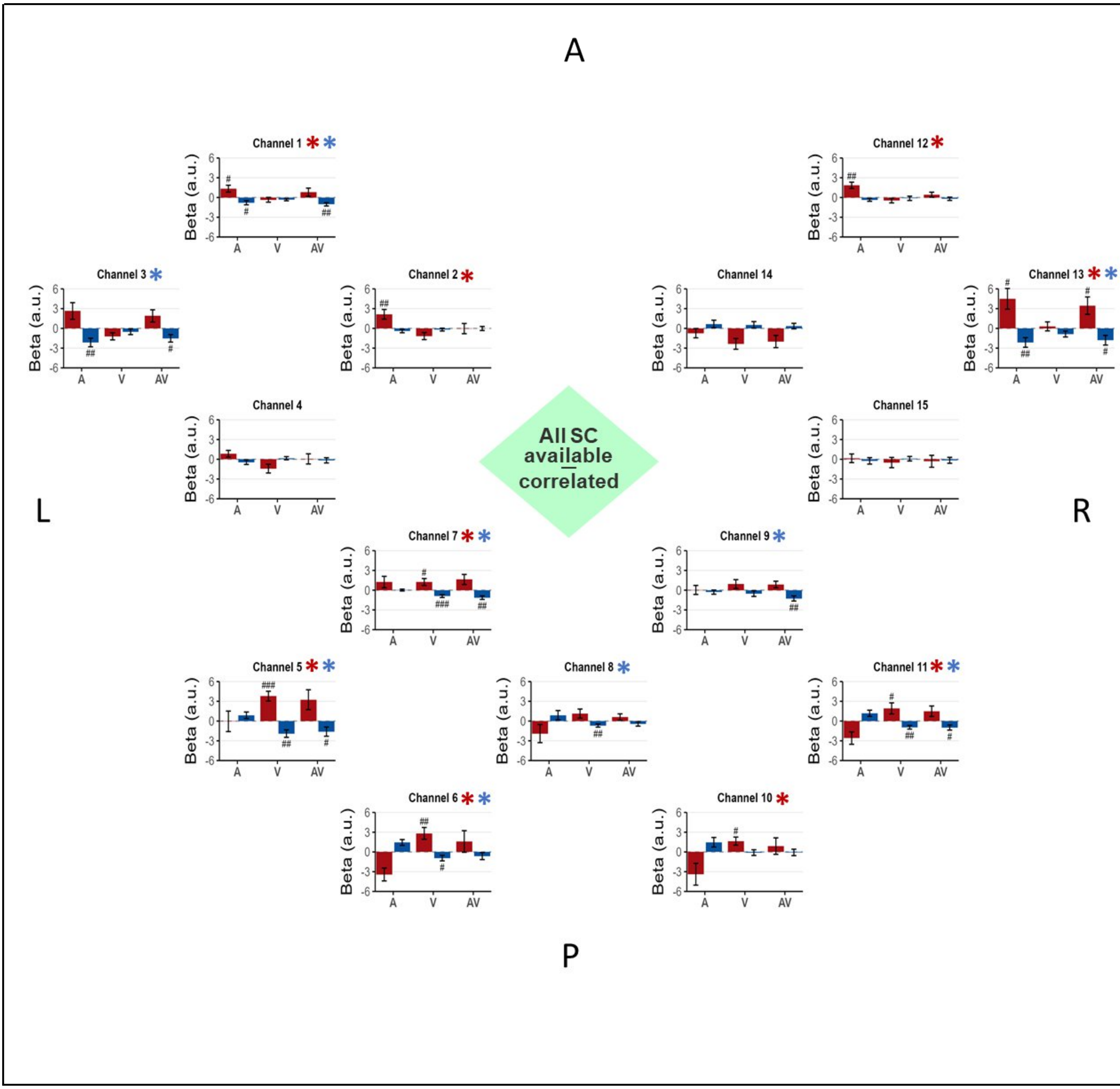


**<u>Figure S5:</u> Full short-channel availability - signal-specific approach (most correlated)**. *Same as in Figure S1, but showing results from the analysis using the most correlated SC available for regression in the full short-channel availability scenario.*

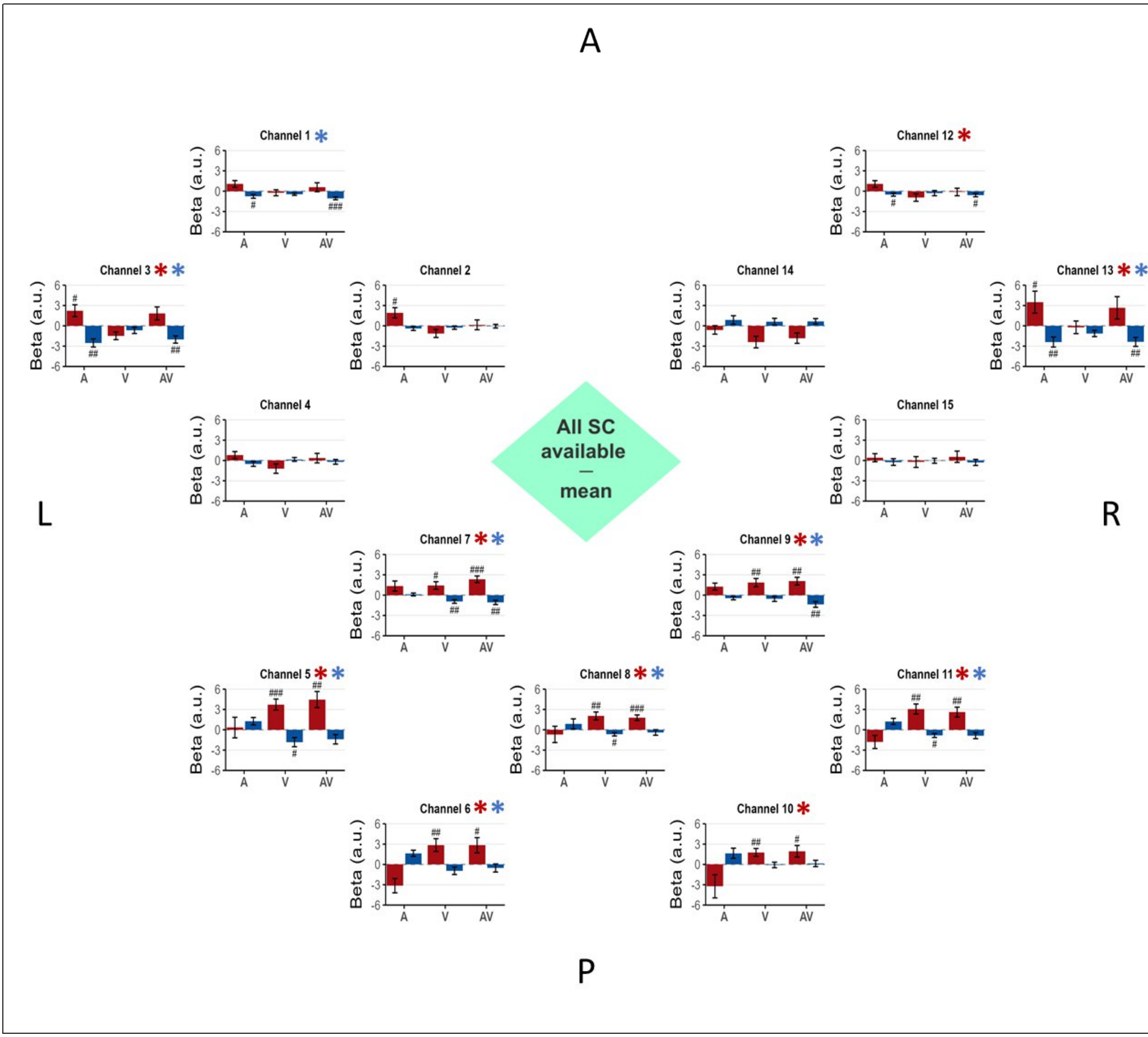


**<u>Figure S6:</u> Full short-channel availability - non-selective mean-based approach**. *Same as in Figure S1, but showing results from the analysis using the mean of all SCs available for regression in the full short-channel availability scenario.*

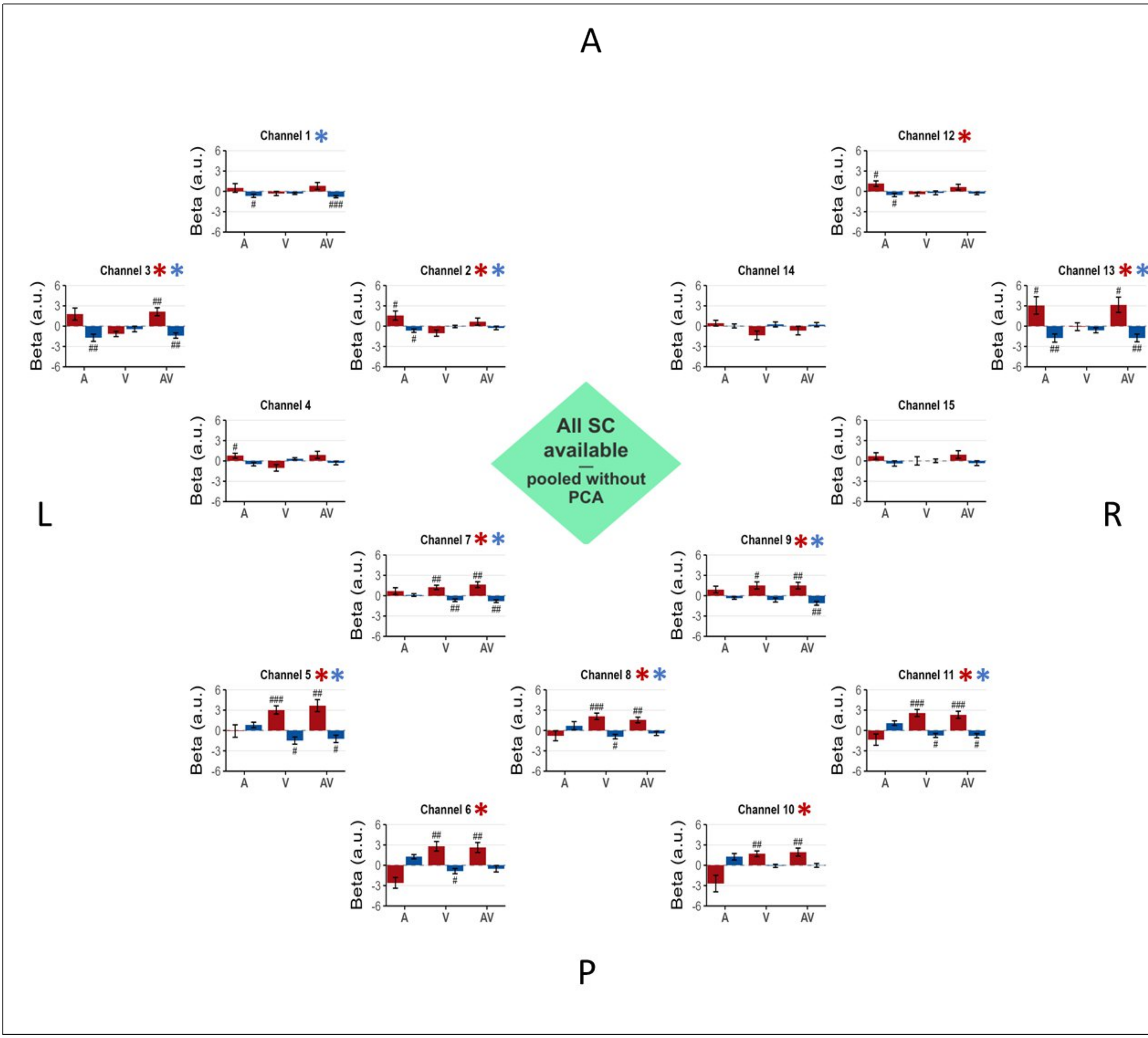


**<u>Figure S7:</u> Full short-channel availability – non-selective approach without PCA**. *Same as in Figure S1, but for the scenario with all short-channel availability. The analysis used raw signal of all available short-channel signals for regression.*
